# CIS checkpoint deletion enhances the fitness of cord blood derived natural killer cells transduced with a chimeric antigen receptor

**DOI:** 10.1101/2020.03.29.014472

**Authors:** May Daher, Rafet Basar, Elif Gokdemir, Natalia Baran, Nadima Uprety, Ana Karen Nunez Cortes, Mayela Mendt, Lucila Nassif Kerbauy, Pinaki P. Banerjee, Mayra Hernandez Sanabria, Nobuhiko Imahashi, Li Li, Francesca Lorraine Wei Inng Lim, Mohsen Fathi, Ali Rezvan, Vakul Mohanty, Yifei Shen, Hila Shaim, Junjun Lu, Gonca Ozcan, Emily Ensley, Mecit Kaplan, Vandana Nandivada, Mustafa Bdaiwi, Sunil Acharya, Yuanxin Xi, Xinhai Wan, Duncan Mak, Enli Liu, Sonny Ang, Luis Muniz-Feliciano, Ye Li, Jing Wang, Shahram Kordasti, Nedyalko Petrov, Navin Varadarajan, David Marin, Lorenzo Brunetti, Richard J. Skinner, Shangrong Lyu, Leiser Silva, Rolf Turk, Mollie S. Schubert, Garrett R. Rettig, Matthew S. McNeill, Gavin Kurgan, Mark A. Behlke, Heng Li, Natalie W. Fowlkes, Ken Chen, Marina Konopleva, Richard Champlin, Elizabeth J. Shpall, Katayoun Rezvani

**Affiliations:** Department of Stem Cell Transplantation and Cellular Therapy, The University of Texas MD Anderson Cancer Center, Houston, TX, USA; Department of Leukemia, The University of Texas MD Anderson Cancer Center, Houston, TX, USA; Department of Stem Cell Transplantation and Cellular Therapy, Hospital Israelita Albert Einstein, Sao Paulo, Brazil; Human Genome and Stem Cell Research Center, Department of Genetics and Evolutionary Biology, Biosciences Institute, University of Sao Paulo, Sao Paulo, Brazil; Department of Chemical and Biomolecular Engineering, University of Houston, Houston, TX, USA; Department of Bioinformatics and Computational Biology, The University of Texas MD Anderson Cancer Center, Houston, TX, USA; System Cancer Immunology, Comprehensive Cancer Centre, King’s College London, London, UK; Center for Cell and Gene Therapy, Baylor College of Medicine, Houston, TX, USA; C.T. Bauer College of Business, University of Houston, Houston, TX, USA; Integrated DNA Technologies, Coralville, IA, USA; Dana-Farber/Harvard Cancer Center, Boston, MA, USA.; Veterinary Medicine & Surgery, The University of Texas MD Anderson Cancer Center, Houston, TX, USA

## Abstract

Immune checkpoint therapy has produced remarkable improvements in the outcome for certain cancers. To broaden the clinical impact of checkpoint targeting, we devised a strategy that couples targeting of the cytokine-inducible SH2-containing (CIS) protein, a key negative regulator of interleukin (IL)-15 signaling, with chimeric antigen receptor (CAR) engineering of natural killer (NK) cells. This combined strategy boosted NK cell effector function through enhancing the Akt/mTORC1 axis and c-MYC signaling, resulting in increased aerobic glycolysis. When tested in a lymphoma mouse model, this combined approach improved NK cell anti-tumor activity more than either alteration alone, eradicating lymphoma xenografts without signs of any measurable toxicity. We conclude that combining CIS checkpoint deletion with CAR engineering promotes the metabolic fitness of NK cells in an otherwise suppressive tumor microenvironment. This approach, together with the prolonged survival afforded by CAR modification, represents a promising milestone in the development of the next generation of NK cells for cancer immunotherapy.

## Introduction

Adoptive cellular therapy using autologous T cells transduced with a chimeric antigen receptor (CAR) has proved to be a powerful approach in the treatment of human cancers, especially B cell leukemias and lymphomas.^1,2^ However, ongoing efforts to consolidate and extend these gains face a number of obstacles: (i) uncoupling cytotoxicity against tumor cells from systemic toxicity, (ii) finding ways to reduce target antigen negative relapses, (iii) overcoming the inhibitory effects of checkpoint molecules in the infused immune effector cells, and (iv) developing universal off-the-shelf cell therapy products that avoid the logistic hurdles of generating autologous products, as well as several of the pitfalls of allogeneic T-cell therapy, such as graft-versus-host disease (GvHD).^3^

Natural killer (NK) cells are attractive candidates for the next wave of effective cancer immunotherapies. They mediate potent cytotoxicity against a range of tumor cells^4^ and, unlike T cells, lack the capacity to induce GvHD in the allogeneic setting.^5^ Moreover, their ready availability in high numbers from various sources, such as umbilical cord blood (CB), boosts their potential as an off-the-shelf product for widespread clinical scalability.^6,7^ One of the most intriguing recent advances in the development of NK cell-based immunotherapy was the demonstration that genetic modification of these cells to express a CAR can enhance their effector function.^8^ This led to the realization that one might overcome some of the limitations of NK cell immunotherapy in cancer, such as the lack of antigen specificity and poor persistence, by exploiting current genetic engineering tools. In our experience, transducing NK cells with a retroviral vector encoding a CD19-specific CAR, the interleukin (IL)-15 cytokine and an inducible suicide element (iCaspase9), can induce greater *in vivo* expansion and longer-term persistence than seen with non-transduced control NK cells. While our preclinical study^9^ using an aggressive model of Raji lymphoma confirmed that this approach can prolong survival times of mice, it was not curative, leading us to question whether the anti-tumor activity of CAR-transduced NK cells could be further enhanced by inhibiting key immune checkpoints in NK cells. Immune checkpoint-based therapies, which target the regulatory pathways of immunocompetent cells to enhance anti-tumor responses, have been at the heart of many recent clinical advances and have led to long-term remissions and possible cures.^10,11^ Most of this success was achieved with T-cells, but there are compelling reasons to predict that checkpoint ablation could modify NK cells in ways that would facilitate their proliferation, persistence and antitumor activity.^12^

The suppressor-of-cytokine signaling (SOCS) family of proteins play important roles in NK cell biology by attenuating cytokine signaling, functional activity, and the immunity of NK cells to cancer.^13,14^ One of its members, the cytokine-inducible SH2-containing protein (CIS), encoded by the *CISH* gene, is induced by cytokines such as IL-2, IL-3 and IL-15^15,16^ and was recently identified as an important intracellular checkpoint molecule in NK cells.^17^ Given that our CAR19-specific NK cells are designed to secrete IL-15, we hypothesized that CIS would be a logical checkpoint molecule to target to enhance the potency and anti-tumor activity of NK cells at lower doses. Here, we show that a combined strategy of CAR engineering and *CISH* knock-out (KO) in CB-derived NK cells results in superior tumor control compared to either approach alone. This gain of effector function is attributed largely to enhanced IL-15 signaling secondary to *CISH* KO with consequent activation of Akt, mTORC1, and c-MYC, leading to increased glycolysis in NK cells in response to tumor targets. Thus, we demonstrate for the first time, that silencing a critical checkpoint in NK cells improves their immune and metabolic ‘fitness’, permitting greater proliferative expansion, persistence and cytotoxic effector function than seen with unmodified NK cells. Our data support the merging of CAR-engineering and immune checkpoint gene editing to enhance the therapeutic potential of NK cells in the clinic.

## Results

### Phenotypic and molecular signaling alterations associated with *CISH* deletion

The central hypothesis of this study predicts that knocking out the *CISH* gene in CAR-transduced NK cells will enhance their effector function against tumor cells, much in the way that targeting PD-1 improves outcomes by removing a critical immune checkpoint in T cells.^18,19^ Our approach for combined retroviral transduction with the iC9/CAR19/IL-15 construct and Cas9 ribonucleoprotein (Cas9 RNP)-mediated gene editing to silence *CISH* is shown in **Fig. 1a,b**. On day 7, we nucleofected CAR-transduced NK cells with Cas9 alone (Cas9 mock) or Cas9 pre-loaded with gRNA targeting *CISH* exon 4. The iC9/CAR19/IL-15 transduction efficiency and cell viability on day 7 were greater than 90% and remained stable over time in both control and gene edited cells (**Supplementary Fig. 1a-b**). The efficiency of *CISH* KO was high (81-98%) in both the non-transduced (NT) control and CAR-expressing NK cells by PCR (**Fig. 1b**) and western blot analysis (**Fig. 1c**) and remained stable over time (**Supplementary Fig. 1c)**. These on target efficiency was confirmed by Sanger sequencing (**Fig. 1d**). Importantly, analysis of *CISH* expression in NK cells either transduced with iC9/CAR19/IL-15 or not transduced (NT control) and then cultured with IL-2 together with K562 feeder cells engineered to express membrane-bound IL-21, 4-1BB ligand and CD48 (henceforth referred to as uAPC) showed a significant time-dependent increase in *CISH* expression in both the CAR-modified and NT control NK cells (**Fig. 1e**). This indicates that CAR-modified and NT-NK cells are subject to the same counter-regulatory circuits leading to modulation of CIS levels. We attribute the higher level of CIS expression in CAR vs. NT-NK cells to the stimulatory effect of IL-15 expressed from the retroviral CAR vector.

**Fig. 1.**
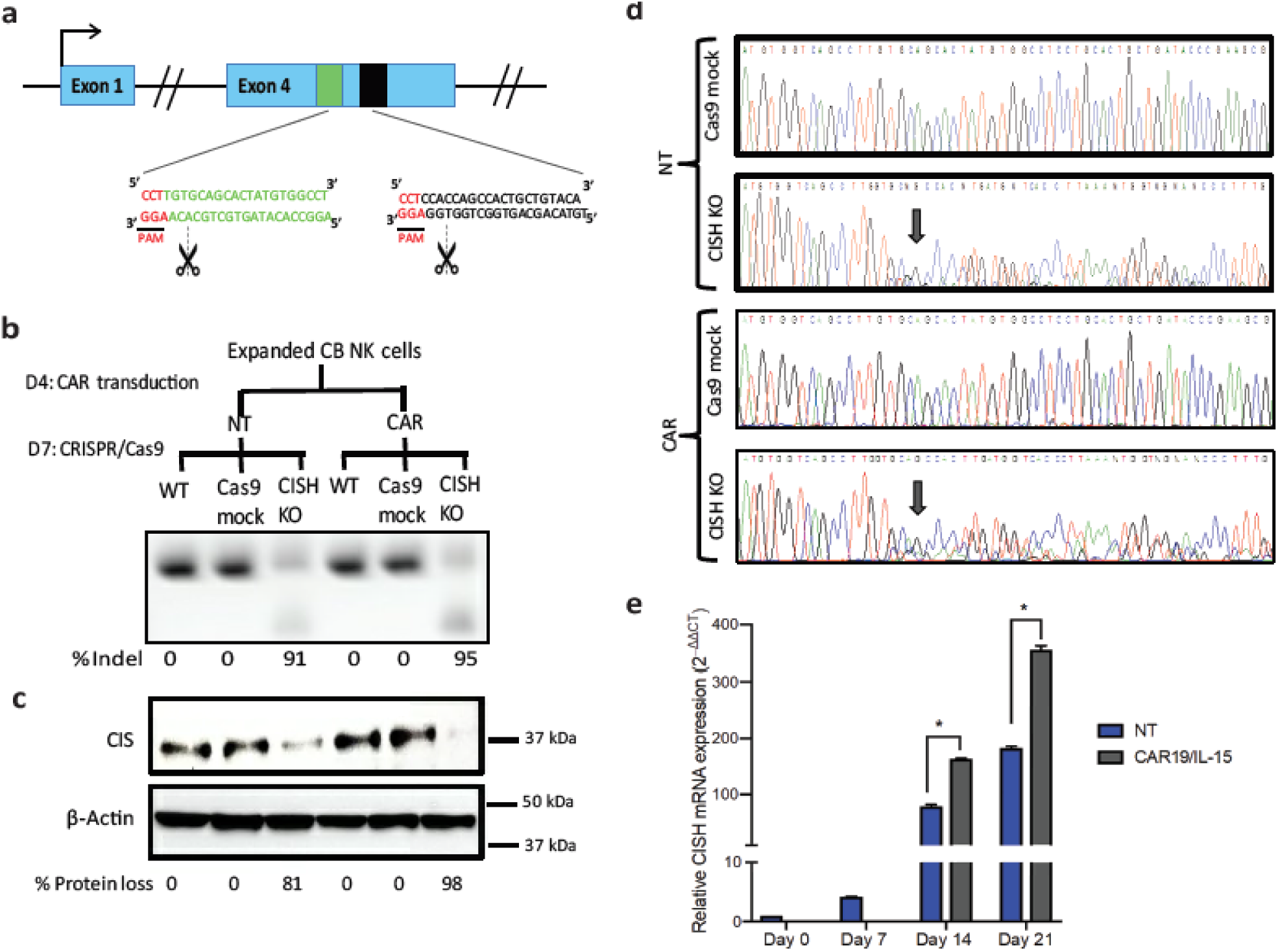
CRISPR/Cas9-mediated deletion of *CISH* in iC9/CAR19/IL-15 NK cells. **a**, Schematic representation of CRISPR/Cas9-mediated *CISH* KO using two guide RNAs (gRNA) targeting exon 4 of the *CISH* gene. **b,c,** CB-NK cells were expanded with K562 based feeder cells and IL-2, then either left non-transduced (NT) or transduced with a retroviral vector expressing iC9/CAR19/IL-15 construct on day 4 of expansion. On day 7 of expansion, NT and iC9/CAR19/IL-15-expressing CB-NK cells were nucleofected with Cas9 alone (Cas9 mock), Cas9 preloaded with gRNA targeting *CISH* exon 4 (*CISH* KO) or non-nucleofected (WT). The *CISH* KO efficiency was determined by PCR (**b**) and western blot analysis (**c**). **d**, Sanger sequencing results showing multiple peaks reflecting non-homologous end joining (NHEJ) events in NT or iC9/CAR19/IL-15 (CAR) NK cells that underwent *CISH* KO compared to single peaks in CTRL (Cas9 mock). Arrows indicate the base pair position where the gene editing started. **e**, Bar graphs showing the relative mRNA expression levels of *CISH* determined on days 0, 7, 14 and 21 of expansion in NT (blue) and iC9/CAR19/IL-15 transduced (grey) NK cells by reverse transcription polymerase chain reaction (RT-PCR) (n=3). Note that on days 0 and 7 only data for NT-NK cells are included since the CAR transduction step is performed on day 4 of expansion.18 S ribosomal RNA (18S) was used as the internal reference gene. Bars represent mean values with standard deviation, *p ≤ 0.05.

Next, to gain insight into the phenotypic changes that accompany *CISH* KO in CAR-transduced NK cells, we used cytometry by time-of-flight (CyToF) and a panel of 37 antibodies against inhibitory and activating receptors, as well as differentiation, homing and activation markers (**Supplementary Table 1**). As shown in **Fig. 2a**, *CISH* KO induced a phenotype characterized by the increased expression of markers of activation and cytotoxicity. These included granzyme-B, perforin, TRAIL and CD3ζ; transcription factors such as eomesodermin (eomes) and T-bet; adaptor molecules such as DAP12; and activating co-receptors/proliferation markers such as DNAM, CD25 and Ki67. A similar profile of upregulated markers was identified after *CISH* KO in NT-NK cells (**Supplementary Fig. 2a**). Using viSNE, a t-distributed stochastic neighbor-embedding (tSNE) algorithm, we further analyzed these NK cells after *CISH* KO, observing marked phenotypic differences between the control and *CISH* KO iC9/CAR19/IL-15 NK cells, again with a predominantly activated phenotype driven by proliferative and cytotoxic features (e.g. increased expression of CD25, Ki67, CD3ζ, perforin and granzyme-b) in *CISH* KO iC9/CAR19/IL-15 NK cells (**Fig. 2b**).

**Fig. 2.**
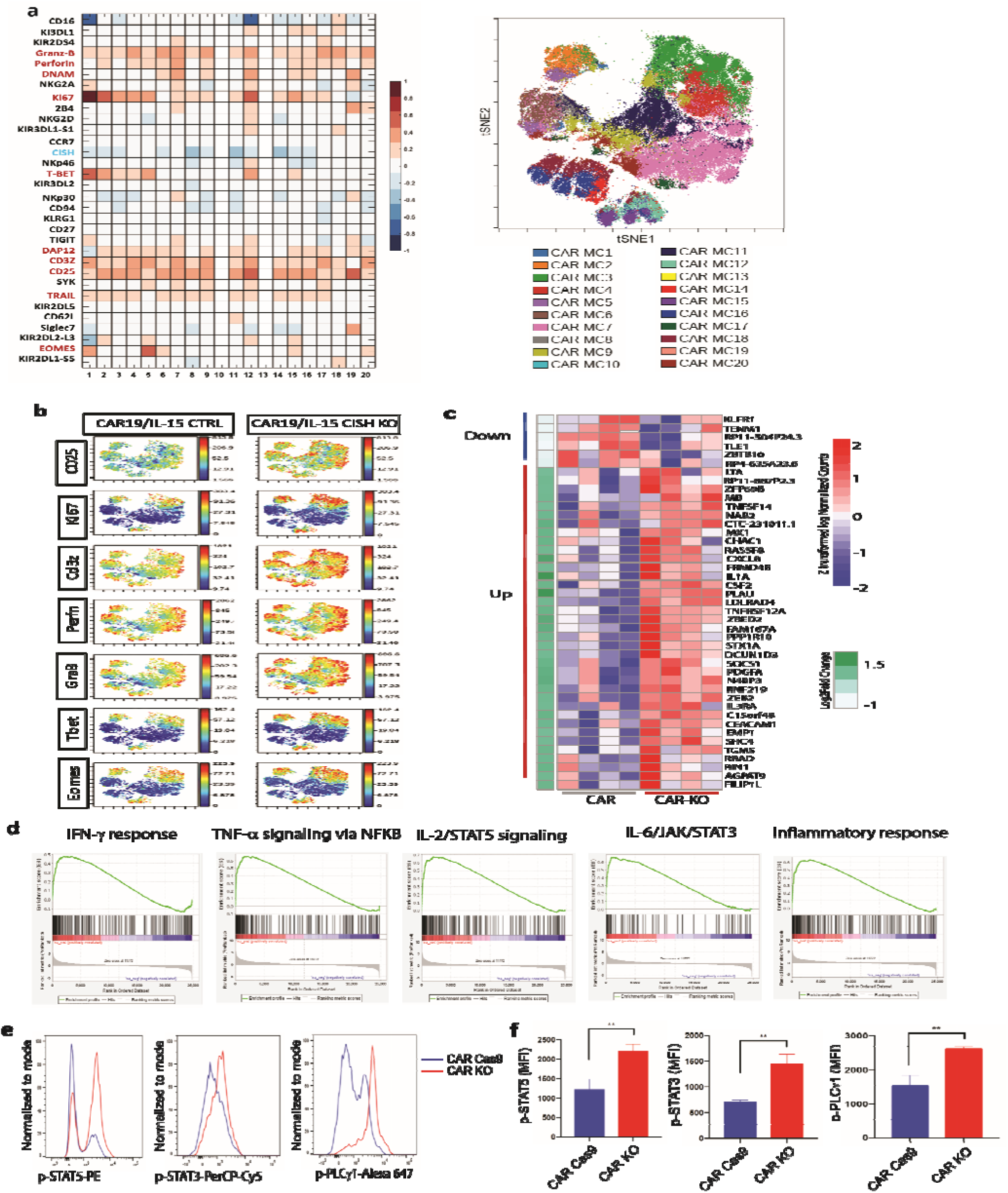
Phenotype and molecular signature of iC9/CAR19/IL-15 *CISH* KO NK cells. **a**, Comparative heatmap of mass cytometry data showing the expression of NK cell surface markers, transcription factors and cytotoxicity markers in iC9/CAR19/IL-15 *CISH* KO compared to iC9/CAR19/IL-15 CTRL NK cells. Each column represents a separate cluster identified by FlowSOM analysis and each row reflects the expression of a certain marker for each annotation. Color scale shows the expression level for each marker, with red representing higher expression and blue lower expression in iC9/CAR19/IL-15 *CISH* KO NK cells. The t-SNE map generated from FlowSOM analysis in the right panel shows the 20 NK cell meclusters (MC) represented in the mass cytometry heat map in the left panel. **b**, Individual t-SNE maps show the expression of selected NK cell markers for iC9/CAR19/IL-15 *CISH* KO compared to iC9/CAR19/IL-15 CTRL NK cells. Color scale indicates signal intensity, ranging from low (blue) to high (red) after arcsine transformation. **c**, Global gene expression analysis by RNA sequencing. Heat map displays the genes that were differentially expressed in purified iC9/CAR19/IL-15 *CISH* KO vs. iC9/CAR19/IL15 CTRL NK cells (n=4). Color scale shows the expression level of each marker, with red representing higher expression and blue lower expression in iC9/CAR19/IL-15 CTRL (CAR) or iC9/CAR19/IL-15 *CISH* KO (CAR KO) NK cells (q< 0.1 and absolute log2foldchange > 0.8). **d,** Gene set enrichment analysis (GSEA) showing enrichment analysis (GSEA) showing enrichment in IFN-γ response, TNF-α signaling via NF-kB, IL-2/STAT5 signaling, IL-6/JAK/STAT3 signaling and inflammatory response in iC9/CAR19/IL-15 *CISH* KO compared to iC9/CAR19/IL-15 CTRL NK cells. **e**, Representative histogram showing enhanced phosphorylation of STAT5 (p-STAT5), STAT3 (p-STAT3) and phospholipase C gamma 1 (p-PLCγ1) in iC9/CAR19/IL-15 *CISH* KO vs. iC9/CAR19/IL-15 CTRL NK cells after co-culture with Raji cells for 30 minutes. Blue histograms represent CAR Cas9 CTRL and red histograms represent CAR *CISH* KO. **f**, Bar graphs showing mean fluorescence intensity (MFI) of p-STAT5, p-STAT3 and p-PLCγ1 in iC9/CAR19/IL-15 *CISH* KO vs. iC9/CAR19/IL-15 CTRL NK cells (n=3). Bars represent mean values with standard deviation, **p ≤ 0.01.

To assess the consequences of *CISH* KO on the transcriptomic and signaling pathway responses of NK cells, we performed RNA sequencing studies of *CISH* KO NT and iC9/CAR19/IL-15 NK cells vs their unmodified controls. *CISH* KO led to upregulation of genes related to inflammatory and immune responses (eg. TNF and IFN signaling such as TNFRSF12A, MX1 and T-bet regulation such as ZEB2) as well as cytokine signaling (eg. IL1A, IL3RA, CXCL8) (**Fig. 2c and Supplementary Fig. 2b**) in both NT and iC9/CAR19/IL-15 NK cells. Thus, we used gene set enrichment analysis (GSEA) to identify sets of genes and biological pathways that could validate and expand on the *CISH* KO phenotype in NT and iC9/CAR19/IL-15 NK cells. This analysis revealed enrichment of genes involved in TNF-α, IFN-γ, IL-2/STAT5 and IL-6/JAK/STAT3 signaling, as well as in those related to inflammatory immune responses (**Fig. 2d and Supplementary Fig. 2c**). We confirmed our findings at the protein level by showing enhanced phosphorylation of STAT5, STAT3 and phospholipase C gamma 1 (PLCγ1) in *CISH* KO iC9/CAR19/Il-15 NK cells (**Fig. 2e,f**). Considered together, these phenotypic and molecular signaling results support the hypothesis that targeting the SOCS family protein CIS in these CAR-modified NK cells removes an important immune checkpoint.

### *CISH* ablation enhances the anti-tumor activity of iC9/CAR19/Il-15 NK cells

When tested against CD19-expressing Raji lymphoma cells, our *CISH* KO iC9/CAR19/IL-15 NK cells produced higher amounts of IFN-γ and TNF-α, displayed greater degranulation (CD107a) and exerted more potent cytotoxicity against their targets than did their respective NT and iC9/CAR19/IL-15-transduced NK controls (**Fig. 3a-d**), consistent with removal of the CIS checkpoint. This outcome was reinforced by the effect of *CISH* KO on the formation of the immunologic synapse (IS) between NT or iC9/CAR19/IL-15 NK cells and Raji cells. Indeed, polarization of perforin-centroid defined microtubule-organizing center (MTOC) was augmented by *CISH* deletion as reflected by a shortened MTOC-to-IS distance compared to controls (**Fig. 3e,f**), a finding that typically correlates with increased effector cell function and cytotoxicity.^20^

**Fig. 3.**
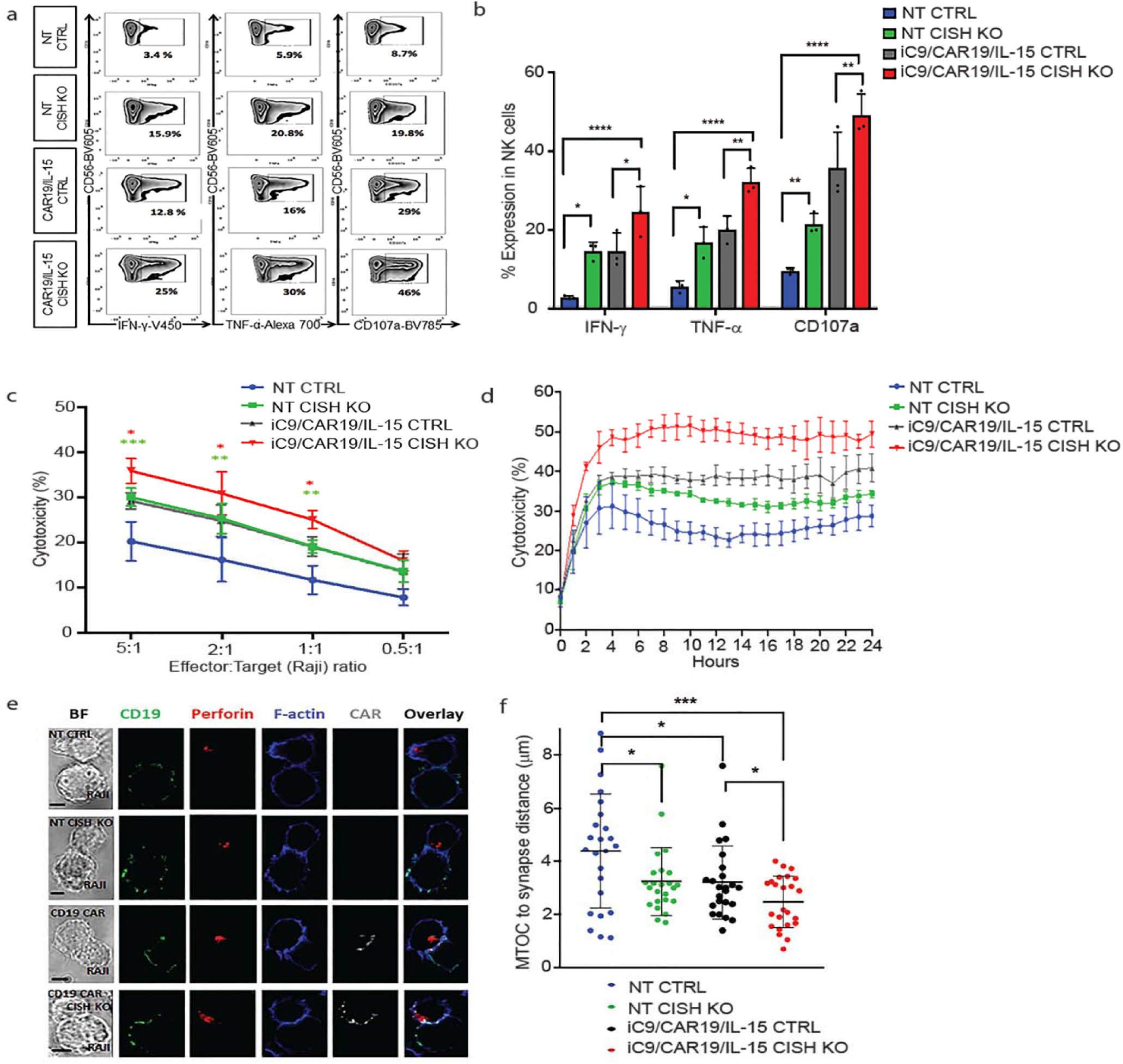
*CISH* deletion improves function and cytotoxicity of NT and iC9/CAR19/IL-15 NK cells. **a,** Representative FACS plots of cytokine production (IFN-γ, TNF-α) and CD107a degranulation by NT CTRL, NT *CISH* KO, iC9/CAR19/IL-15 CTRL or iC9/CAR19/IL-15 *CISH* KO NK cells after co-culture with Raji target cells for 6 hours. Inset values indicate the frequency of IFN-γ-, TNF-α- and CD107a-positive cells from each group. **b,** Bar plots summarize the flow cytometry data on cytokine production (IFN-γ and TNF-α) and CD107a degranulation by NT CTRL, NT *CISH* KO, iC9/CAR19/IL-15 CTRL or iC9/CAR19/IL-15 *CISH* KO NK cells after co-culture with Raji target cells for 6 hours (n= 3). Statistical significance is indicated as *p≤ 0.05, ** p ≤ 0.01, ****p ≤ 0.0001, bars represent mean values with standard deviation. **c**, Cytotoxicity of NT CTRL, NT *CISH* KO, iC9/CAR19/IL-15 CTRL or iC9/CAR19/IL-15 *CISH* KO NK cells against Raji targets at different effector: target (E:T) ratios, as measured by ^51^Cr-release assay (n=3). The bars represent mean values with standard deviation. The red asterisks represent the statistical significance between iC9/CAR19/IL-15 *CISH* KO vs. iC9/CAR19/IL-15 CTRL NK cells, *p≤ 0.05. The green asterisks represent the statistical significance between NT *CISH* KO vs. NT CTRL NK cells, ***p ≤ 0.001, ** p ≤ 0.01. **d**, Cytotoxicity of NT CTRL, NT *CISH* KO, iC9/CAR19/IL-15 CTRL or iC9/CAR19/IL-15 *CISH* KO NK cells against Raji targets over 24 hours at 1:1 E:T ratio as measured by Incucyte live imaging cell killing assay (n=3). Bars represent mean values with standard deviation. At 10 hrs: NT *CISH* KO vs. NT CTRL (p=0.04), iC9/CAR19/IL-15 *CISH* KO vs iC9/CAR19/IL-15 CTRL (p=0.007). **e**, Confocal microscopy showing representative synapse images of NT CTRL, NT *CISH* KO, iC9/CAR19/IL-15 CTRL or iC9/CAR19/IL-15 *CISH* KO NK cells conjugated with Raji cells. Images show conjugates in bright field (BF) or stained with anti-CD19 (green), anti-perforin (red), phalloidin-F-actin (blue), anti-CAR (grey) and an overlay of fluorescence channels are also shown. **f**, NT CTRL, NT *CISH* KO, iC9/CAR19/IL-15 CTRL or iC9/CAR19/IL-15 *CISH* KO NK cells were assessed for their ability to polarize lytic granules to Raji cells as measured by distance from microtubule-organizing center (MTOC) to the immune synapse. Results from 3 independent donors are shown. Each data point represents a single immunologic synapse. Statistical significance is indicated as *p≤ 0.05, ***p ≤ 0.001.

### Robust metabolic changes associated with CIS checkpoint elimination

Our decision to ablate the CIS immune checkpoint as a therapeutic strategy was partly based on evidence that this protein functions as a critical negative regulator of IL-15 signaling in NK cells.^17,21^ Thus, by releasing a potent brake on IL-15 activity, we sought to enhance NK cell activation, expansion and cytotoxic function over extended times. Although this approach is supported by *in vitro* studies,^17,21^ we remained concerned over the possible impact of increased IL-15 signaling on prominent metabolic pathways in NK cells. In one negative scenario, long-term exposure to IL-15 could suppress rather than boost metabolic rates, leading to NK cell exhaustion.^22^ In another scenario, it might produce undue systemic toxicity.^23,24^ We therefore evaluated the effects of *CISH* knockout, with or without iC9/CAR19/IL-15 transduction, on two major regulators of NK cell metabolism: mTORC1, which controls pathways responsible for proliferation and cytotoxicity,^25,26^ and MYC, which upregulates glucose transporters and glycolytic enzymes that promote glycolysis.^27^ Indeed, GSEA analysis revealed an enrichment of genes involved in PI3K/Akt/mTOR, mTORC1, c-MYC and glycolysis (**Fig. 4a**).

**Fig. 4.**
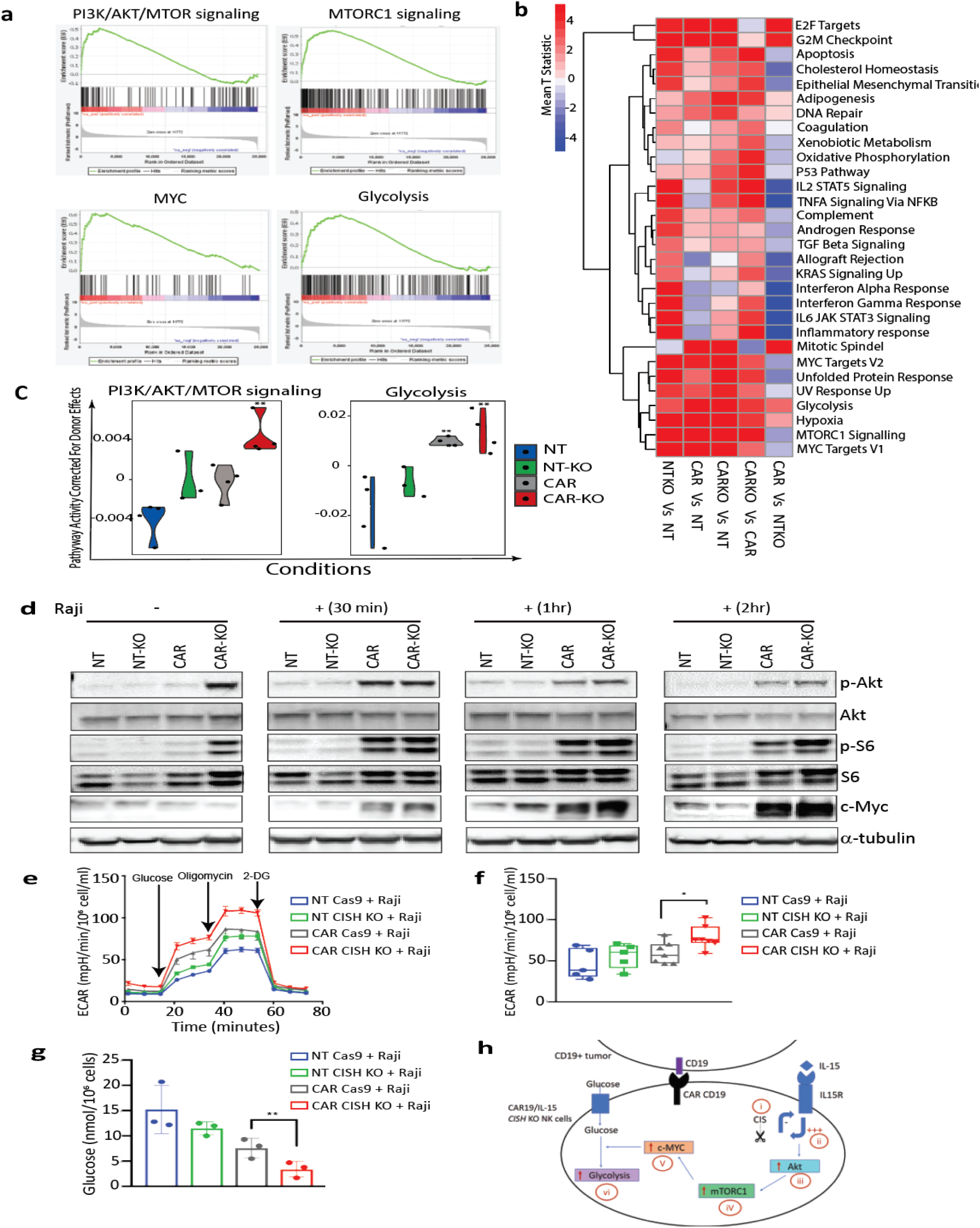
Metabolic changes associated with iC9/CAR19/IL-15 *CISH* KO NK cells. **a**, GSEA plots showing enrichment in PI3K/Akt/MTOR, mTORC1, MYC and glycolysis pathways in iC9/CAR19/IL-15 *CISH* KO NK cells compared to iC9/CAR19/IL15 CTRL NK cells. **b**, Comparative mean T statistic heat map of RNA sequencing data showing the expression of metabolic pathways in NT CTRL (NT), NT *CISH* KO (NT-KO), iC9/CAR19/IL-15 CTRL (CAR) or iC9/CAR19/IL-15 *CISH* KO (CAR-KO) NK cells that are significantly different (q < 0.01) in at least one of the five comparisons. Each column represents a separate comparison and each row reflects the expression of a certain hallmark pathway for each annotation. Color scale indicates signal intensity, ranging from lower (blue) to higher (red) expression. **c**, Violin plots showing PI3K/Akt/mTORC1 and glycolysis signaling in NT (blue), NT-KO (green), CAR (grey) or CAR-KO (red) NK cells after correction for donor effect. Pathway activity of samples is regressed against donor and the residual is the corrected pathway activity. p-values reported are computed relative to NT using the linear regression approach discussed in the methods. **p≤ 0.01. **d**, NT CTRL (NT), NT *CISH* KO (NT-KO), iC9/CAR19/IL-15 CTRL (CAR) or iC9/CAR19/IL-15 *CISH* KO (CAR-KO) NK cells were cultured without (-) or with (+) Raji cells for 30 min, 1h or 2h, NK cells were then purified and the protein expression levels of p-Akt, Akt, p-S6, S6, c-MYC and α-tubulin in NK cells were determined by western blot analysis. Representative blots from two independent experiments are shown. **e**, A series of extracellular acidification rate (ECAR) were calculated for NT CTRL (blue lines), NT *CISH* KO (green lines), iC9/CAR19/IL-15 CTRL (grey lines) or iC9/CAR19/IL-15 *CISH* KO (red lines) NK cells co-cultured with Raji targets for 2hrs and subsequently purified and treated with 2 g/L D-glucose, 1 μM oligomycin and 100 mM 2-Deoxyglucose (2-DG). A representative graph is shown from five independent experiments. **f**, Box plots summarize the ECAR data by NT CTRL (blue box), NT *CISH* KO (green box), iC9/CAR19/IL-15 CTRL (grey box) or iC9/CAR19/IL-15 *CISH* KO (red box) NK cells co-cultured with Raji (n=5). Statistical significance is indicated as *p≤ 0.05; bars represent mean values with standard deviation. **g**, Bar graph summarizes the glucose concentration in the supernatant of the different NK cell conditions co-cultured with Raji for 2 hrs: NT CTRL (blue), NT *CISH* KO (green), iC9/CAR19/IL-15 CTRL (grey) or iC9/CAR19/IL-15 *CISH* KO (red) NK cells (n=3). Bars represent mean values with standard deviation, **p≤ 0.01. **h**, Proposed model of the metabolic activity of iC9/CAR19/IL-15 *CISH* KO NK cells. The consequences of *CISH* KO are depicted in red. (i) Silencing *CISH* (ii) enhances IL-15 signaling leading to (iii) increased Akt and (iv) mTORC1 axis; the resulting (v) enhanced c-MYC activation in the presence of tumor renders these cells more metabolically active by (vi) increasing their glycolysis and their ability to respond to immune stimulation.

The above results raise a pivotal question. Are the metabolic gene expression patterns seen with *CISH* KO distinct, or do they overlap with those seen with unmodified or CAR19/IL-15 transduced NK cells? The heatmap in **Fig. 4b** displays the various hallmark pathways and how they change (upregulated or downregulated) among the various comparisons. In general, compared to NT-NK cells, each engineering strategy (*CISH* KO or CAR transduction) alone or in combination led to enrichment of metabolic pathways (**Fig. 4b**). We observed some overlap between the pathways upregulated by *CISH* ablation or iC9/CAR19/IL-15 transduction alone; however, the combination of both approaches clearly led to more robust metabolic changes (**Fig. 4b**). Of note, certain pathways were specifically activated after *CISH* KO and were not induced with CAR transduction alone; these included pathways related to cytokine signaling and inflammatory response (**Fig. 4b**).

We next focused on the metabolic pathways that are functionally relevant for NK cell anti-tumor activity; overall our linear regression model showed an additive effect of *CISH* ablation and CAR transduction, with the largest increase in PI3K/Akt/mTOR and glycolysis pathways being achieved with a combination of the two strategies, suggesting that both genetic manipulations are needed to achieve optimal NK cell effector functions (**Fig. 4c**). This interpretation was confirmed by greater phosphorylation of Akt and ribosomal protein S6 (S6), a downstream target of mTORC1 pathway activation, and by increased expression of c-MYC, an important mediator of glycolysis, in particular in *CISH* KO CAR19/IL-15 NK cells in response to Raji lymphoma cell targets (**Fig. 4d**). Notably, pS6 and pAkt were upregulated in *CISH* KO CAR19/IL-15 NK cells even in the absence of Raji. It is likely by removing the CIS brake, the endogenously secreted IL-15 by CAR19/IL-15 transduced NK cells can trigger the mTORC1/Akt pathway with a lower threshold of activity. In contrast c-MYC, which is the precursor of glycolysis, was only upregulated in response to Raji tumor. To pursue the functional implications of these findings, we first blocked the mTORC1 pathway by treating the CAR19/IL-15 NK cells (+/− *CISH* KO) with rapamycin and observed a decrease in their cytotoxicity against Raji lymphoma compared to untreated cells (**Supplementary Fig. 3a**), which asserts the importance of the mTORC1 pathway as part of the mechanism by which *CISH* KO enhances the cytotoxicity of CAR19/IL15 NK cells. We then used the Seahorse assay to measure the glycolytic response of NK cells to tumor targets and showed that in response to Raji lymphoma cells, either *CISH* KO alone or CAR transduction alone in NK cells, could increase glycolysis, as measured by extracellular acidification rate (ECAR), although the best result (and the only statistically significant one) was achieved by combining *CISH* KO and CAR19/IL-15 transduction (**Fig. 4e,f**). Consistent with these findings, CAR-transduced NK cells with *CISH* deletion showed the greatest glucose consumption, when co-cultured with Raji tumor, compared with controls (**Fig. 4g**). Importantly, in the absence of Raji lymphoma tumor targets, there was no difference in glycolysis among the different NK conditions (**Supplementary Fig. 3b,c**) which parallels the c-MYC expression profile in response to Raji. *CISH* KO iC9/CAR19/IL-15 NK cells also had higher oxygen consumption rate (OCR) compared to control iC9/CAR19/IL-15 NK cells (**Supplemetary Fig 4a**) and induced an increase in the number of mitochondria and the mitochondrial/nuclear volume ratio as assessed by confocal miscroscopy (**Supplementary Fig 4b, c)**. These data suggest that *CISH* KO can also enhance the metabolism of CAR-NK cells by increasing mitochondrial activity.

Finally, we propose a model (**Fig. 4h**), in which *CISH* KO in CAR-transduced NK cells enhances IL-15 signaling by releasing a checkpoint brake, in turn leading to increased activation of the Akt/mTORC1 axis and c-MYC activation, culminating in greater glycolytic capacity of NK cells and a resultant increase in their ability to respond to tumor targets.

### CIS checkpoint disruption combined with iC9/CAR19/IL-15 transduction improves tumor control in a Raji lymphoma model

Using an aggressive Raji lymphoma mouse model (**Fig. 5a**), we next investigated whether adoptive transfer of *CISH* KO NK cells could boost the control of disease in tumor-bearing mice. First, mice received a single intravenous (i.v.) infusion of NK cells (10×10^6^/mouse) that were either unmodified (NT control) or electroporated with Cas9 alone (Cas9 control), or had *CISH* KO. Tumor growth was monitored by changes in tumor bioluminescence imaging (BLI) over time. The tumor burden increased in all animals through day 28 of the study, with no significant differences in survival noted between the *CISH* KO group and controls (**Fig. 5b,c**). We next investigated the *in vivo* antitumor activity of NK cells modified with both *CISH* KO and iC9/CAR19/IL-15 transduction. Because this combination had shown greater potency than either modification alone, even at low E:T ratios *in vitro*, we hypothesized that it would also be more effective at controlling Raji lymphoma cells at lower infusion doses. Indeed, when as few as 3×10^6^ *CISH* KO iC9/CAR19/IL-15 NK cells were administered, they significantly improved survival (p=0.003) and the control of Raji lymphoma compared to results with control NK cells, although the mice eventually succumbed to tumor by day 46 (**Fig. 5d-f**). To improve these results, we infused the animals with a higher dose (10×10^6^) of the *CISH* KO, iC9/CAR19/IL-15 NK cells. This approach eradicated lymphoma in all treated mice as demonstrated by BLI analysis and pathologic examination, and led to significantly prolonged survival times (**Fig. 5d-g; Supplementary Fig. 5a, b and Supplementary Fig. 6)**. This result was associated with improved NK cell persistence (up to 7 weeks after infusion) in mice that received the *CISH* KO iC9/CAR19/IL-15 cells compared to iC9/CAR19/IL-15 NK controls (**Fig. 5h and Supplementary Fig. 5c**).

**Fig. 5.**
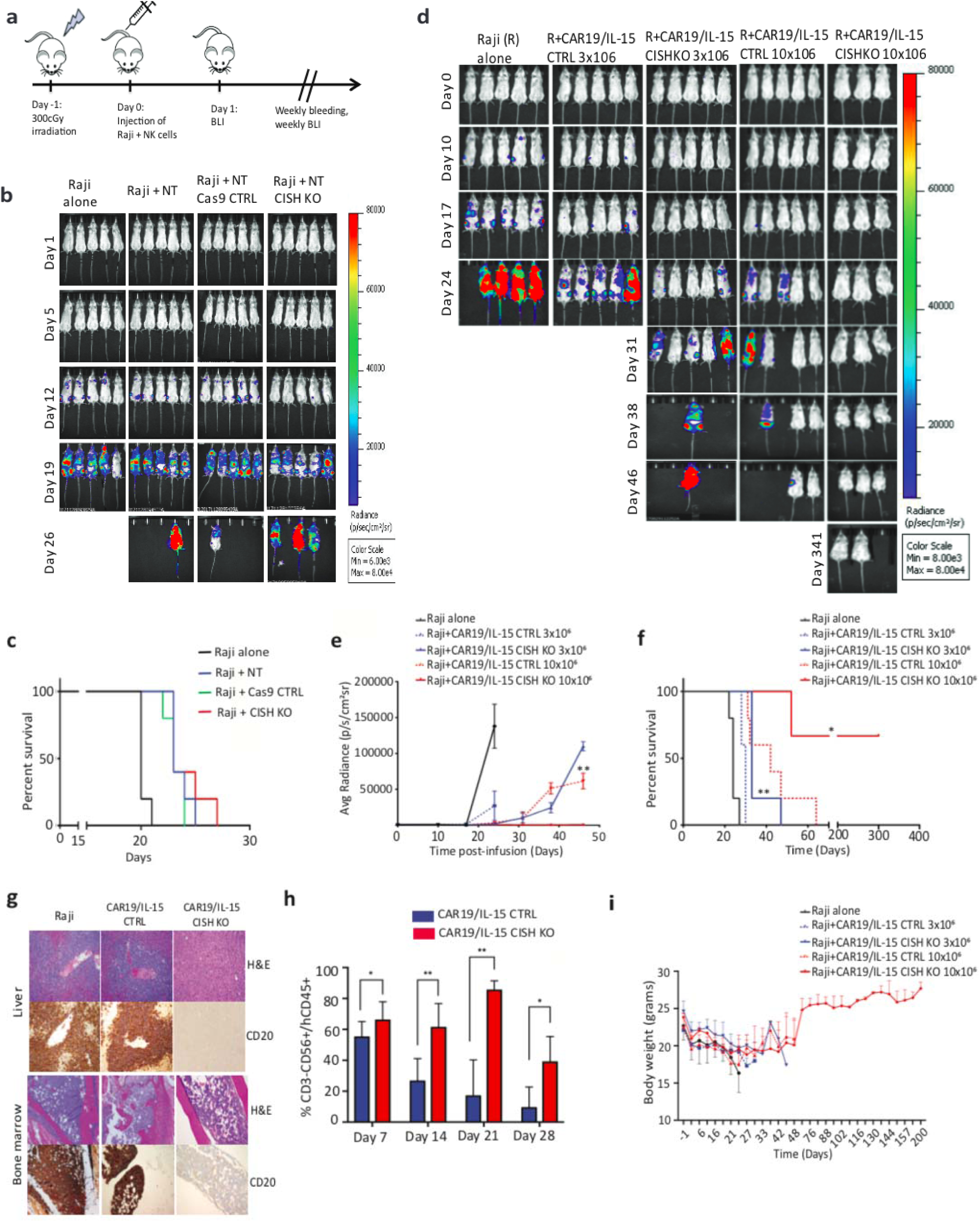
*CISH* KO iC9/CAR19/IL-15 NK cells improve tumor control and survival in a Raji lymphoma mouse model at low infusion doses. **a**, Schematic diagram representing the timeline of the *in vivo* experiments. **b**, Bioluminescent imaging (BLI) was used to monitor the growth of FFluc-labeled Raji tumor cells in NSG mice. The mice were treated with Raji alone, Raji plus one dose of 10 x 10^6^ of NT, NT Cas9 CTRL or NT *CISH* KO NK cells (5 mice per group). Colors indicate intensity of luminescence with red representing higher and blue lower intensity. **c**, Kaplan Meier plots showing the probability of survival for the four groups of mice described in panel b. **d**, BLI data from five groups of NSG mice treated with Raji alone (n=5), Raji plus one dose of 3 x 10^6^ of iC9/CAR19/IL-15 CTRL NK cells (n=5) or iC9/CAR19/IL-15 *CISH* KO NK cells (n=5) or Raji plus one dose of 10 x 10^6^ of iC9/CAR19/IL-15 CTRL NK cells (n=5) or iC9/CAR19/IL-15 *CISH* KO NK cells (n=3). The average radiance (**e**) and survival curves (**f**) are shown for the five groups of mice described in panel d. Statistical significance is represented by *p≤ 0.05 or **p≤ 0.01. **g**, Photomicrographs of H&E and immunohistochemical CD20 staining of liver (top) and bone marrow (BM) (bottom) from mice engrafted with Raji B cell lymphoma and treated with iC9/CAR19/IL-15 *CISH* KO or iC9/CAR19/IL-15 CTLR NK cells (10 x 10^6^ dose level). Representative images show absence of neoplastic B cells in liver and bone marrow of a mouse treated with iC9/CAR19/IL-15 *CISH* KO NK cells in comparison to similar treatment with iC9/CAR19/IL-15 NK cells retaining *CISH* expression. Images were taken at 10x (liver) and 5x (bone marrow) using a Leica DFC 495 camera. **h**, Bar graph showing the percentage of NK cells (CD3-CD56+CD45+) present in peripheral blood from mice treated with iC9/CAR19/IL-15 CTRL vs. iC9/CAR19/IL-15 *CISH* KO NK cells at days 7, 14, 21 and 28. Bars represent mean values with standard deviation. Statistical significance is represented by *p≤ 0.05 or **p≤ 0.01. **i**, Graph showing body weights of NSG mice groups described in panel d over time.

Of note, *CISH* KO was not associated with any signs of increased toxicity in mice, including organ damage or increases in systemic inflammatory cytokines or increased weight loss compared to control groups (**Fig. 5g,i; and Supplementary Fig. 7**). After the initial weight loss observed in all groups after irradiation, mice treated with *CISH* KO iC9/CAR19/IL-15 cells recovered their weight to baseline over an extended period of follow up (**Fig.5i**). Moreover, CAR-transduced NK cells were not detectable in the organs of any animals at autopsy, indicating that *CISH* KO does not induce uncontrolled proliferative expansion and persistence of CAR NK cells. We conclude that silencing of *CISH* in the context of CAR19/IL-15 transduction in NK cells can secure robust control of tumor cells *in vivo* without appreciable toxicity.

### The presence of IL-15 in the CAR construct is necessary for *CISH* KO-mediated improvement in NK cell function

Since our CAR construct expresses both CAR19 and IL-15, it was important to understand whether the improvement in NK cell function following *CISH* KO is dependent on the presence of IL-15 in the CAR construct. To address this question, we transduced NK cells with a retroviral vector expressing CAR19 without the IL-15 transgene, (henceforth referred to as CAR19 NK cells) and analyzed their *CISH* expression after culture with IL-2 and uAPC. CAR19 NK cells showed a time-dependent increase in *CISH* expression to the same extent as observed with NT control, but to a lower magnitude than in iC9/CAR19/IL-15 NK cells (**Supplementary Fig. 8a)**. The efficicency of *CISH* KO in CAR19 NK cells was similar to what was achieved in CAR19/IL-15 NK cells (**Supplementary Fig. 8b**) and led to improvement in their cytotoxicity against Raji tumor targets in vitro (**Supplementary Fig. 8c)** reminescent of the effect we observed in NT NK cells following *CISH* KO. However, when tested in our our Raji lymphoma mouse model, *CISH* KO failed to improve the antitumor activity of CAR19 NK cells in vivo (**Supplementary Fig. 8d-f)**, pointing to the essential role of IL-15 in mediating the effect of *CISH* KO on CAR-NK cell anti-tumor activity.

### Safety evaluation of *CISH* KO iC9/CAR19/IL-15 NK cells for relapsed/refractory B-cell malignancy

To investigate the possibility that enhanced IL-15 signaling may result in autonomous or dysregulated growth of *CISH* KO CB-NK cells, we cultured NT (control or *CISH* KO) or iC9/CAR19/IL-15-transduced CB-NK cells (control or *CISH* KO) in media without the addition of exogenous IL-2 or stimulation with feeder cells for 45 days. Cultured *CISH* KO NT or iC9/CAR.19/IL-15-transduced CB-NK cells did not show any signs of abnormal growth over 6 weeks (**Supplementary Fig. 9**), after which the cells stopped expanding.

Even though *CISH* KO CAR-transduced NK cells lacked evidence of serious toxicity in our *in vivo* model, we planned additional *in vitro* and *vivo* experiments to ascertain that *CISH* KO CAR NK cells could be swiftly eliminated in the event of toxicity in early phase clinical testing. Thus, we relied on the presence of *iC9* as a suicide gene in our vector to confirm that *CISH* KO, CAR-transduced NK cells could be induced to undergo apoptosis in the presence of a small-molecule dimerizer, AP1903. The addition of as little as 10 nM of AP1903 to cultures of CAR-transduced NK cells induced their apoptosis within 4 hours, and *CISH* KO did not affect the action of the dimerizer (**Supplementary Fig. 10a**). The suicide gene was also effective at eliminating the CAR NK cells *in vivo* (**Supplementary Fig. 10b,c**). Mice engrafted with Raji tumor received either control or *CISH* KO CAR-transduced NK cells (n = 5 mice per group) followed by treatment with the dimerizer on days 7 and day 9 after NK cell infusion. The animals were then sacrificed on day 12. Administration of the small-molecule dimerizer resulted in a striking reduction of the transduced cells (both control and *CISH* KO) in the blood and tissues (liver, spleen and bone marrow) in all treated mice (**Supplementary Fig. 10b,c**).

Identifying possible off-target editing events mediated by the CRISPR/Cas9 RNP complexes is crucial before the approach described here can be moved to the clinic. Thus, we used GuideSeq and rhAmpSeq^TM^ technologies (Integrated DNA Technologies) to assess the landscape of genome-wide off-target effects for the specific *CISH* gRNAs tested in our experiments (see materials and methods). GuideSeq experiments were performed using HEK293 cells that constitutively express *S.p* Cas9 nuclease paired with highly-modified synthetic gRNAs. We followed previously published methods^28^ to identify off-target sites with the highest potential to be edited for each crRNA (**Fig. 6a,b**). The potential Cas9 off-target cleavage sites identified by GuideSeq were then quantified in NK cells electroporated with RNP complexes targeting the *CISH* locus using rhAmpSeq technology, a multiplexed targeted enrichment approach for next generation sequencing (NGS). Cells treated with wild type *S.p.* Cas9 protein had a low frequency of off-target editing events with either crRNA1, crRNA2, or the combination of both crRNAs (**Fig. 6c,d**). The use of a high fidelity Cas9 protein (Alt-R HiFi Cas9 v3, IDT)^29^ further reduced the risk of off-target events to <0.5% (**Fig. 6c,d**). These data support the translation of this approach to the clinic.

**Fig. 6.**
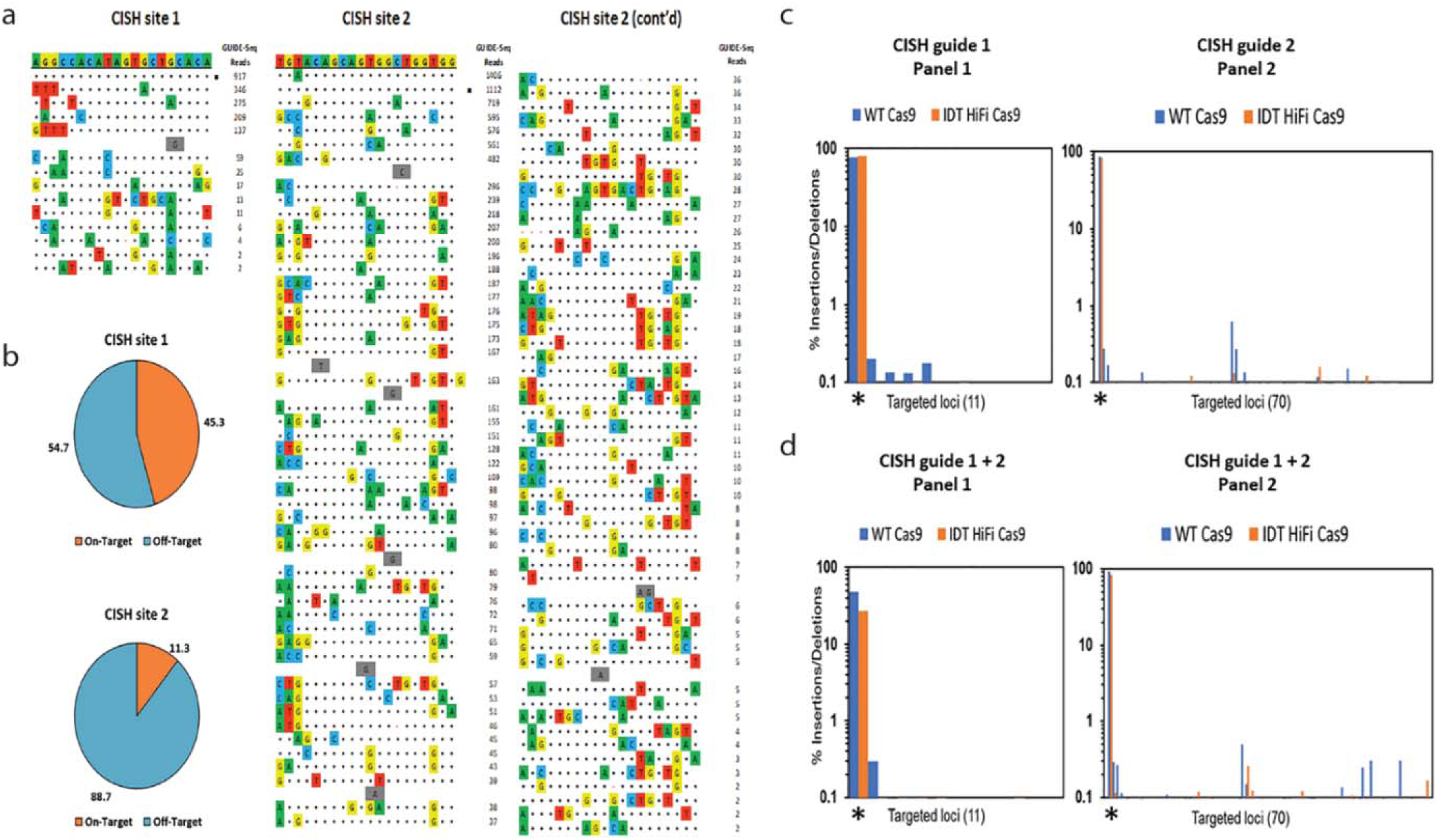
Identification of Cas9 off-target sites by GUIDE-Seq and quantification of potential Cas9 off-target cleavage sites using rhAmpSeq technology. **a,** Sequences of off-target sites identified by GUIDE-Seq for two guides targeting the *CISH* locus. The guide sequence is listed on top with off-target sites shown below. The on-target site is identified with a black square. Mismatches to the guide are shown and highlighted in color with insertions shown in grey. The number of GUIDE-Seq sequencing reads are shown to the right of each site. 10 μM Alt-R crRNA XT complexed to Alt-R tracrRNA was delivered into HEK293 cells that constitutively express Cas9 nuclease by nucleofection. **b,** Pie charts indicate the fractional percentage of the total unique, CRISPR/Cas9 specific read counts that are on-target (orange) and off-target (blue). Total editing at the on- and off-target sites identified by GUIDE-Seq was measured using rhAmpSeq, a multiplexed targeted enrichment approach for next generation sequencing (NGS). For each of the two *CISH* targeting guides amplicons were designed around each Cas9 cleavage site with reads >1% of the on target GUIDE-Seq reads. RNP complexes formed with either WT Cas9 (blue) or Alt-R HiFi Cas9 (orange) were delivered via electroporation into expanded NK cells. **c,** INDEL formation at each targeted loci for *CISH* guide 1 (panel 1, 11-plex) and *CISH* guide 2 (panel 2, 70-plex) when a single RNP complex was delivered. The on-target locus is indicated with a black asterisk underneath the first 2 bars of each graph. **d,** INDEL formation at each targeted loci when *CISH* guide 1 and *CISH* guide 2 were co-delivered. The on-target locus is indicated with a black asterisk underneath the first 2 bars of each graph.

## Discussion

It is now widely appreciated that NK cell differentiation, effector function and survival, defined as ‘fitness’, are coupled to metabolic reprogramming processes. However, signals and checkpoints that regulate NK cell fitness and function in the tumor microenvironment are not well defined. For instance, the relevance of the classical T-cell checkpoints PD-1 and CTLA-4 in NK cell the biology is not clear. Even after activation, CTLA-4 is not expressed in NK cells.^30^ On the other hand, PD-1 has been reported to be expressed on memory-like NK cells after viral infections^31^ while others correlate its upregulation on NK cells with lower anti-tumor activity.^32–34^ Our study focuses on a more defined NK specific checkpoint molecule, CIS, which has been shown to play an important role as a negative regulator of cytokine signaling in murine NK cells.^17^

Having demonstrated the enhanced expansion, persistence and antitumor activity of CB-derived NK cells engineered to express IL-15 and a CD19-redirected CAR^9^, we hypothesized that by combining this modification with removal of a vital cytokine checkpoint, it might be possible to boost NK cell immune fitness still farther. Here we report that combining CAR- and IL-15-transduced NK cells with *CISH* deletion to remove the IL-15 cytokine checkpoint enhances antitumor activity more than either strategy alone. This effect was especially striking in a mouse xenograft model of Raji lymphoma, where the combination approach uniformly eradicated tumor xenografts (at the 10×10^6^ dose level), while infusion of the singly modified NK cells proved inadequate. Moreover, the doubly modified NK cells persisted nearly twice as long as control CAR19/IL-15 transduced NK cells. We attribute this gain of effector function to IL-15-driven Akt/mTORC1 and MYC signaling, to removal of the CIS checkpoint, with a consequent increase in glycolytic activity. These findings have clinical relevance because they establish, to our knowledge for the first time, proof-of-principle that targeting a critical cytokine checkpoint in NK cells will augment the benefits of constitutive CAR expression, thus promoting the immune fitness of these lymphocyte subsets.

Previous reports have shown that *CISH* deletion in NK cells is beneficial in tumor models only where IL-15 was present in the tumor microenvironment.^17,21^ In fact, in a multiple myeloma mouse model, where IL-15 is a minor cytokine in the malignant milieu, *CISH* deletion in NK cells did not alter the natural history of the cancer or the survival of the mice.^21^ In our study in xenografted mice, there was no appreciable increase in the activity of non-transduced NK cells or CAR19 transduced NK cells (without IL-15 transgene) after *CISH* KO; the only improvement was noted in animals treated with *CISH* KO CAR19/IL15-transduced NK cells. This suggests that the robust anti-tumor activity of NK cell with *CISH* KO depends upon the availability of IL-15 in the tumor microenvironment. Although the systemic delivery of IL-15 has considerable appeal, this strategy has been limited by severe toxicity.^23,24^ Thus, incorporation of the *IL-15* gene in the CAR construct, which is not associated with the toxicities seen with systemic IL-15, likely played an important role in the enhanced anti-tumor activity of *CISH* KO iC9/CAR19/IL-15 NK cells in our tumor model. Given the absence or limited concentration of IL-15 in the microenvironment of some cancers, expressing this cytokine in the CAR construct is essential to achieving an optimal therapeutic effect from *CISH* knockout.

Our results suggest that removal of the CIS checkpoint may increase the metabolic fitness of NK cells by enhancing the activity of downstream metabolic pathways. In the model we propose (**Fig. 4h**), *CISH* KO releases the brake on IL-15 signaling, which in turn enhances Akt/mTORC1 activity^26^, leading to the upregulation of c-*MYC*,^35,36^ specifically in response to tumor targets. The expression of c-MYC is regulated by the T cell antigen receptor (TCR) and by pro-inflammatory cytokines such as interleukin-2 (IL-2) in T-cells and by IL-15 in NK cells.^37,38^ We have previously shown that upon activation, CAR19/IL-15 NK cells secrete more IL-15.^9^ We now propose that upon activation by the tumor, the higher levels of IL-15 in the microenvironment leads to increased IL-15/Akt/mTORC1 and c-MYC signaling in *CISH* KO CAR19/IL-15 NK cells. Furthermore, c-MYC controls expression of the metabolic machinery in NK cells, specifically a switch to aerobic glycolysis (Warburg effect).^39,40^ This accelerated glycolytic flux leads to an accumulation of glycolytic intermediates, which are channeled into biosynthetic pathways to generate the DNA, lipids and proteins needed for an effective anti-tumor immune response.^41^ We would emphasize that in our studies, the shift in metabolism towards aerobic glycolysis was not observed in *CISH* KO iC9/CAR19/IL-15 CB-derived NK cells in the absence of tumor and was only apparent when the NK cells were cultured with their tumor targets, which parallels the c-MYC expression profile and supports the hypothesis postulated above. This observation is important, as an exaggerated increase in glycolysis in *ex vivo* expanded T cells has been reported to severely impair the ability of CD8+ T-cells to persist long-term and form memory cells *in vivo*.^42^ Hence, careful tuning of NK cell metabolism to ensure a timely increase in aerobic glycolysis in response to tumor stimulation would be desirable; however, further research is needed to elucidate the immunometabolism of CAR-NK cells *in vivo*, especially in microenvironments such as solid tumors, where nutrients and oxygen are restricted.

In summary, this is the first report of a genetic engineering strategy combining CAR transduction and *CISH* deletion in CB-derived NK cells. When tested in a preclinical tumor model, this cellular product eliminated CD19+ lymphoma cells even at relatively low doses, without the induction of serious toxicity. Our findings support the merging of CAR-engineering and cytokine checkpoint gene editing to enhance the therapeutic potential of NK cells in the clinic.

## Materials and Methods

### Cell lines and culture media

The Raji cell line (Burkitt lymphoma cell line) was purchased from American Type Culture Collection (Manassa, VA, USA). K562 based feeder cells were retrovirally transduced to co-express 4-1BBL, CD48 and membrane bound IL-21 (referred to in the text as uAPC). All cell lines were cultured in Roswell Park Memorial Institute (RPMI) medium supplemented with 5% fetal bovine serum (FBS), 1% penicillin-streptomycin and 1% L-glutamine. NK cells were cultured in Stem Cell Growth Medium (SCGM) supplemented with 5% FBS, 1% penicillin-streptomycin and 1% L-glutamine.

### Cord blood NK cell expansion

CB units for research were provided by the MD Anderson CB Bank. Lymphocytes were isolated by a density-gradient technique (Ficoll-Histopaque; Sigma, St Louis, MO, USA). CD56+ NK cells, purified with an NK isolation kit (Miltenyi Biotec, Inc., San Diego, CA, USA), were then stimulated with irradiated (100 Gy) uAPC (K562 based feeder cells that are retrovirally transduced to co-express 4-1BBL, CD48 and membrane bound IL-21) at 2:1 feeder cell:NK ratio and recombinant human IL-2 (Proleukin, 200 Chiron, Emeryville, CA, USA) in complete stem cell growth medium (SCGM, CellGenix GmbH, Freiburg, Germany) on day 0. Activated NK cells were transduced with retroviral supernatants on day +4 in human fibronectin-coated plates (Clontech Laboratories, Inc., Mountain View, CA, USA). On day +7 and day +14, NK cells were stimulated again with irradiated feeder cells and IL-2CAR-transduced NK cells were collected for use on day 21of culture.

### Retrovirus transfection and transduction

The retroviral vector encoding iC9.CAR19.CD28-zeta-2A-IL-15 has been previously described^43,44^ and was kindly provided by Dr. Gianpietro Dotti (University of North Carolina). The retroviral vector encoding CAR19.CD28-zeta (without IL-15) was used as a control in some of the experiments where indicated. Transient retroviral supernatants were produced as previously described.^44^ Activated NK cells were purified and transduced with retroviral supernatants on day +4 in human fibronectin-coated plates (Clontech Laboratories, Inc., Mountain View, CA, USA). Three days later (day +7), CAR transduction efficiency was measured by flow cytometry and NK cells were stimulated again with irradiated feeder cells and IL-2.

### CRISPR/Cas9 gene editing of *CISH*

*CISH* KO was performed on day +7 using ribonucleoprotein (RNP) complex, in both NT and CAR transduced NK cells. Protospacer sequences for the *CISH* gene were identified using the CRISPRscan algorithm (www.crisprscan.org).^45^ DNA templates for gRNAs were made using the protocol described by Li et al.^46^ We used two gRNAs targeting exon 4 of the human *CISH* gene: crRNA1:AGGCCACATAGTGCTGCACA, crRNA2: TGTACAGCAGTGGCTGGTGG. Cas9 protein (PNA bio) and gRNA were incubated at room temperature for 15 min in a 3:1 reaction with 15μg Cas9 and 5000 ng gRNA (2500ng of sgRNA#1 and 2500ng of sgRNA#2).^47^ The incubation product (gRNA and Cas9 complex) was then used to electroporate 1-2 million NT or CAR-transduced NK cells using the Neon transfection system (Thermo Fisher Scientific). Alt-R HiFi Cas9 Nuclease 3-NLS (IDT) was also used in the above-described protocol to form RNP complexes with both gRNAs for electroporation into NT or CAR-transduced NK cells. Optimized electroporation conditions were 1600 V, 10 ms, 3 pulses, using T buffer. The different cell preparations were then co-cultured with irradiated uAPC at a 1:2 ratio (NK:uAPC) in SCGM media with IL-2 200 U/ml. For the off-target studies of sgRNA#1 and sgRNA#2, Alt-R crRNA and ATTO-labeled tracRNA (IDT) comprised the guide RNA; RNP complexes were formed with both Alt-R WT and Alt-R HiFi S.p. Cas9 and electroporated into NK cells.

### Real-time quantitative PCR (RT Q-PCR)

Total RNA was isolated using the RNeasy Plus Mini kit (Qiagen Inc., Hilden, Germany) and cDNA synthesis performed with ReadyScript cDNA Synthesis Mix (Sigma-Aldrich Corp., St. Louis, MO, USA) according to the manufacturer’s instructions. PCR reaction mix of 20 μL of TaqMan 2X Advanced Fast PCR MasterMix (Applied Biosystems Inc., Foster City, CA, USA), 1 μL of CISH TaqMan Gene Expression Assay (Applied Biosystems), 2 μL of cDNA and 7 μL of nuclease-free water. Primer sequences and PCR conditions have been described.^48^ Real-time Q-PCR was performed on an ABI 7500 Fast Real Time PCR System (Applied Biosystems). mRNA levels were quantified against standard curves generated using sequential dilutions of an oligonucleotide corresponding to each amplified PCR fragment and using 7500 Fast v2.3 Software (Applied Biosystems). Relative expression was determined by normalizing the amount of each gene of interest to the housekeeping gene 18S.

### Western blot

To detect CIS expression, NK cells were pre-treated with 10 μM MG132 for 4 h to block proteasomal degradation. NK cells were then lysed in lysis buffer (IP Lysis Buffer, Pierce Biotechnology Inc., Rockford, IL) supplemented with protease inhibitors (Complete Mini, EDTA-free Cocktail tablets, Roche Holding, Basel, Switzerland) and incubated for 30 min on ice. Protein concentrations were determined by the BCA assay (Pierce Biotechnology Inc., Rockford, IL). The following primary antibodies were used: CIS antibody (Clone D4D9) and β-actin antibody (Clone 8H10D10), both antibodies were obtained from Cell Signaling Technology. For the western blots performed to assess the metabolic changes, the same protocol was used except that MG132 was only added for the last 30 min (or during NK cell selection for the samples co-cultured with Raji) and during lysis. NK cells were purified before lysis using NK isolation kit (Miltenyi Biotec, Inc., San Diego, CA, USA). The following primary antibodies were used: phospho-S6 ribosomal protein antibody (Ser240/244) (Clone D68F8) and S6 ribosomal protein (Clone 5G10) antibody, p-Akt (Ser473) (Clone D9E) and total Akt (clone 4604) and c-Myc antibody (Clone 9E10) were obtained from Cell Signaling Technology. Alfa-tubulin antibody was purchased from Abcam (Clone DM1A). Blots were imaged using the LI-COR Odyssey Infrared Imaging System.

### Assessing knockout efficiency using PCR gel electrophoresis

DNA was extracted and purified (QIAamp DNA Blood Mini Kit, Qiagen Inc., Hilden, Germany) from CAR-transduced and *ex vivo*-expanded NT NK cells (control and *CISH* KO conditions). We used the Platinum™ SuperFi™ Green PCR Master Mix from Invitrogen for PCR amplification using the following PCR primers spanning the Cas9-sgRNA cleavage site of the exon 4 of *CISH* gene:

Exon 4 Forward Primer: CGTCTGGACTCCAACTGCTT
Exon 4 Reverse Primer: GTACAAAGGGCTGCACCAGT

DNA bands were separated by polyacrylamide gel electrophoresis prepared with SYBR safe DNA gel stain in 0.5xTBE. Gel images were obtained using GBox machine with GeneSys software (Syngene, Frederick, MD). Band intensities were analyzed using imageJ software by plotting band intensities of each lane. % cleavage was calculated by the ratio of the intensities of the cleaved bands to uncleaved bands.

### Sanger sequencing to measuring allele modification frequencies

DNA was extracted and purified (QIAamp DNA Blood Mini Kit, Qiagen Inc., Hilden, Germany) from CAR-transduced and *ex vivo*-expanded NT NK cells (control and *CISH* KO conditions). The sequences of the PCR primers used for the amplification of the target locus are provided in the PCR gel electrophoresis section. The purified PCR products were sent for Sanger Sequencing at MD Anderson’s core facility using both PCR primers, and each sequence chromatogram was analyzed with the online Finch TV software available at: http://www.geospiza.com/Products/finchtv.shtml

### NK cell functional assays

On days 14 or day 21 of culture, control *ex vivo*-expanded NT (control or *CISH* KO) and CAR19/Il-15 NK cells (control or *CISH* KO) at 100 × 10^3^ cells/well were co-cultured in the presence of Brefeldin A for 6uh in round bottom 96-well plates with Raji cells or K562 targets (positive control) at an effector:target cell ratio (E:T) of 2:1. Cells were then collected and washed with PBS and surface and intracellular staining performed using BD fixation/permeabilization kit. To be able to gate on NK cells the following surface antibodies were used: Live/dead aqua dead cell stain (Thermofisher), APC/Cy7 anti-human CD3 (HIT3a, Biolegend), Brilliant Violet 605™ anti-human CD56 (NCAM) Antibody (HCD56, Biolegend). CAR transduction was measured using anti-CAR antibody Goat F(ab’)2 anti-Human IgG (H+L)-Alexa Fluor 647 (Jackson Laboratories). CD107a degranulation was measured with anti-human CD107a (LAMP-1) conjugated to Brilliant Violet 785™, Biolegend, San Diego, CA, USA) added to the wells at the beginning of co-culture. Intracellular cytokine production was determined using antibodies against TNFα (MAb11) conjugated to Alexa Fluor 700, eBioscience Inc., San Diego, CA, USA) and against IFN-γ conjugated to V450 (BD Biosciences, San Jose, CA, USA) by flow cytometry as previously described.^49^

### Phospho flow cytometry assays

iC9/CAR19/IL-15 *CISH* KO and iC9/CAR19/IL-15 CTRL NK cells were co-cultured in 96 well plates with Raji cells at 1:1 ratio (10^5^ NK cells and 10^5^ Raji cells per well x 3 wells) for 30 minutes. Cells were then collected, washed and phosphoflow staining was done using the Perfix Expose Kit from Beckman Coulter (B26976) per manufacturer’s instructions. To be able to gate on NK cells the following surface antibodies were used: Live/dead aqua dead cell stain (Thermofisher), APC/Cy7 anti-human CD3 (HIT3a, Biolegend), Brilliant Violet 605™ anti-human CD56 (NCAM) Antibody (HCD56, Biolegend). The following antibodies were used for intracellular staining: BD Phosflow™ PerCP-Cy™5.5 Mouse anti-Stat3 (pY705), BD Phosflow™ PE Mouse Anti-Stat5 (pY694) (BD biosciences), BD Alexa Fluor® 647 Mouse Anti-Human PLCγ (pY783). The following isotype control antibodies were used: PerCP-Cy™5.5 Mouse IgG2a, κ Isotype Control, PE Mouse IgG1, κ Isotype Control, Alexa Fluor® 647 Mouse IgG1 κ Isotype control, all purchased from BD biosciences.

### Cytotoxicity assays

#### Chromium release assay

To assess cytotoxicity, *ex vivo* expanded NT NK cells (control and *CISH* KO) and CAR-transduced (control and *CISH* KO) were co-cultured with ^51^Cr-labeled Raji targets at multiple E:T ratios; cytotoxicity was measured by ^51^Cr release as previously described.^49^

#### Incucyte real-time cytotoxicity assay

*Ex vivo* expanded NT NK cells (control and CISH KO) and CAR-transduced (control and CISH KO) were co-cultured in 96 well well plates at 1:1 E:T ratio with Raji tumor targets pre-labeled with Vybrant DyeCycle Ruby Stain (ThermoFisher) and CellTracker Deep Red Dye (ThermoFisher). Each well contained 5 x10^4^ Raji cells and 5 x 10^4^ NK cells. Apoptosis was detected using the CellEvent Caspase-3/7 Green Detection Reagent (ThermoFisher). Frames were captured over a period of 24hrs at 1-hour intervals from 4 separate 1.75 x 1.29 mm^2^ regions per well with a 10× objective using IncuCyte S3 live-cell analysis system (Sartorius). Values from all four regions of each well were pooled and averaged across all three replicates. Results were expressed graphically as percent cytotoxicity by calculating the ratio of overlapping red and green signal (counts per image) divided by red signal (counts per image). Where indicated, NK cells were pre-incubated with Rapamycin (100ng/ml) for 4hrs prior to initiation of the incucyte assay.

### Confocal microscopy for studying the immunologic synapse

NK cells (0.5×10^6^) were conjugated with 0.25×10^6^ Raji cells in 250μl of SCGM with 10% heat inactivated FBS containing media, for 40 minutes at 37^0^C and stained as described previously.^9,50^ Briefly, after incubation, cells were adhered onto a Poly-A-Lysine coated slide (Electron Microscopy Sciences) and stained for proteins of interest. Alexa Fluor 647-conjugated affinity-purified F(ab’)2 fragment goat anti-human IgG (H+L) antibody was used to detect CAR. Anti-CD19 Alexa Fluor (AF) 488 (clone HIB19, BD BioSciences), Phalloidin AF 568 (Invitrogen) for detection of F-actin, and anti-Perforin 421 (clone δ G9; BioLegend) were used. Conjugates were mounted in anti-fade containing media (Prolong gold, Invitrogen) and were imaged by sequential scanning with a Yokogawa spinning disk confocal microscope equipped with a Zyla 4.2sCMOS Camera, and under 63x objective. Images were exported to Imaris (Bitplane) for quantitative measurements. The distance from perforin centroid to synapse was measured as previously described.^50,51^

### Mass Cytometry

#### Antibody conjugation

A panel comprising of 37 metal-tagged antibodies was used for the in-depth characterization of NK cells (Table S1). All unlabeled antibodies were purchased in carrier-free form and conjugated in-house with the corresponding metal tag using Maxpar X8 polymer per manufacturer’s instructions (Fludigm). All metal isotopes were acquired from Fludigm except for indium (III) chloride (Sigma-Aldrich, St. Louis, MO). Antibody concentration was determined by measuring the amount of A280 protein using Nanodrop 2000 (Thermo Fisher Scientific, Waltham, MA). Conjugated antibodies were then diluted in PBS-based antibody stabilization solution or LowCross-Buffer (Candor Bioscience GmbH, Wangen, Germany) supplemented with 0.05% sodium azide (Sigma-Aldrich, St. Louis, MO, USA) to a final concentration of 0.5 mg/ml. Serial titration experiments were performed to determine the concentration with the optimal signal-to-noise ratio for each antibody.

#### Sample preparation, staining and acquisition

NT-NK cells (control or *CISH* KO) and CAR transduced NK cells (control or *CISH* KO) were harvested, washed twice with cell staining buffer (0.5% bovine serum albumin/PBS) and incubated with 5 μl of human Fc receptor blocking solution (Trustain FcX, Biolegend, San Diego, CA) for 10 minutes at room temperature. Per previously published protocol^52^, cells were then stained with a freshly prepared antibody mix against cell surface markers for 30 minutes at room temperature on a shaker (100 rpm). For the last 3 minutes of incubation, cells were incubated with 2.5 μM cisplatin (Sigma Aldrich, St Louis, MO), washed twice with cell staining buffer and fixed/permeabilized using BD Cytofix/Cytoperm solution for 30 minutes in the dark at 4°C. Cells were washed twice with perm/wash buffer, stained with antibodies directed against intracellular markers and after an additional wash step, stored overnight in 500 μl of 1.6% paraformaldehyde (EMD Biosciences)/PBS with 125 nM iridium nucleic acid intercalator (Fluidigm). The next day, samples were washed twice with cell staining buffer, resuspended in 1 ml of MilliQ dH2O, filtered through a 35 μm nylon mesh (cell strainer cap tubes, BD, San Jose, CA) and counted. Before analysis, samples were suspended in MilliQ dH2O supplemented with EQTM four element calibration beads at a concentration of 0.5×105/ml. Samples were acquired at 300 events/second on a Helios instrument (Fluidigm) using the Helios 6.5.358 acquisition software (Fluidigm).

#### Data analysis

Mass cytometry data were normalized based on EQ^TM^ four element signal shift over time using Fluidigm normalization software 2. Initial data quality control was performed using Flowjo version 10.4.1. Calibration beads were gated out and singlets were chosen based on iridium 193 staining and event length. Dead cells were excluded by the Pt195 channel and further gating was performed to select CD45+ cells and then the NK cell population of interest (CD3-CD56+). A total of 320,000 cells were proportionally sampled from all samples to perform automated clustering. Data were analyzed using automated dimension reduction including (viSNE) in combination with FlowSOM for clustering^53^ for the deep phenotyping of immune cells as published before^54^. We further delineated relevant cell clusters, using our in-house pipeline for cell clustering (CytoClustr (published^55^ and available upon request))

### RNA sequencing

RNA was extracted and purified with the RNeasy Plus Mini Kit (Qiagen) from CAR-transduced and *ex vivo*-expanded NT NK cells (control and *CISH* KO conditions) and sent for RNA sequencing at MD Anderson’s core facility, where quality control, library construction and sequencing were performed. Analysis of RNAseq data was performed by the MD Anderson Bioinformatics Department. The Toil RNA-seq workflow^56^ was used to convert RNA sequencing data into gene- and transcript-level expression quantification. CutAdapt was used to remove extraneous adapters, STAR was used for alignment and read coverage, and RSEM were used to produce quantification data. DEseq2^57^ was used to carry out pairwise differential expression between groups of interest. The model also included the donor of the sample to control for donor specific effects. The T-statistic generated by DESeq2 was used to carry out GSEA analysis using the gage package^58^. Differentially expressed pathways were identified at a q < 0.01. Barcode plots for pathways of interest were generated using the broad institutes GSEA tool kit (http://software.broadinstitute.org/gsea/index.jsp).

ssGSEA analysis^59^ was performed using the GSVA package^60^. To infer the effect of each condition (NTKO, CAR and CARKO) on pathway activity relative to NT cells we made use of the following linear regression models. E ∼ Donor+Condition: Where E is pathway activity and Donor was the donor individual that the sample was obtained from. A p-value and t-statistic were generated for each condition relative to NT. Pathways of interest are pathways where effect size in CARKO > CAR/NTKO relative to NT. (significant pathways are identified at p < 0.01). For both GSEA and ssGSEA analysis we used the Hallmark geneset.^61^

### Seahorse extracellular acidification assays

Extracellular acidification rate (ECAR) and oxygen consumption rate (OCR) were measured using the Agilent Seahorse XFe96 Analyzer (Agilent) as per the manufacturer’s instructions. NT NK cells (Control or CISH KO) and CAR transduced NK cells (Control or *CISH* KO) alone or after culturing with Raji, were previously washed twice in PBS and resuspended in glycolysis stress test media: DMEM media supplemented with 2 mM L-Glutamine, prewarmed to 37° and pH adjusted to 7.4. Next cells were plated in a 96-well plate, precoated with Cell Tak, at a density of 400,000 viable cells per well and allowed to attach to the bottom of the plate by gently spinning at 1000 g without break. For measurement of acute cell dependency on glucose and glucose utilization, Glycolysis Stress Test assay was utilized and glucose at a final concentration of 10 mM, oligomycin at 1 μM and 2-deoxyglucose at 100 mM were added during the assay. All assays were performed in 4 or 5 replicates per condition and repeated in 5 independent experiments. Generated data were normalized to viable cell number in million/ml at the viability was found to be consistently between 96% and 99% as determined by trypan blue exclusion assay (ViCell, Beckmann Coulter) and analyzed using Seahorse Glycolysis Stress Test Generator (Agilent).

### Xenogeneic lymphoma models

To assess the antitumor effect of CAR-transduced CB-NK cells *in vivo*, we used a NOD/SCID IL-2Rγnull (NSG) xenograft model, with the aggressive NK-resistant Raji cell line. Mouse experiments were performed in accordance with NIH recommendations under protocols approved by the Institutional Animal Care and Use Committee. NSG mice (10–12 weeks old; Jackson Laboratories, Bar Harbor, ME, USA) were irradiated with 300 cGy at day-1 and inoculated intravenously with firefly luciferase-labeled Raji cells (2 × 10e4) on day 0. Where indicated, fresh expanded NT (control or *CISH* KO) or CAR19/IL15-transduced CB-NK (control or *CISH* KO) or CAR19 (no IL-15)-transduced CB-NK (control or *CISH* KO) cells were injected through the tail vein on day 0. Mice were subjected to weekly bioluminescence imaging (Xenogen-IVIS 200 Imaging system; Caliper, Waltham, MA, USA). Signal quantitation in photons/second was performed by determining the photon flux rate within standardized regions of interest using Living Image software (Caliper).^62^ The mice that died between any two scheduled timepoints for BLI imaging were imaged just prior to sacrifice and the average radiance at the time of death was included in the statistical analysis for the subsequent BLI time point. Trafficking, persistence and expansion of NK cells were measured by flow cytometry. The following antibodies were used for flow cytometry staining: APC/Cy7 anti-human CD3 Antibody (clone HIT3a, Biolegend), Brilliant Violet 605™ anti-human CD56 (NCAM) Antibody (Clone HCD56, Biolegend), anti-CAR antibody Goat F(ab’)2 anti-Human IgG (H+L)-Alexa Fluor 647 (Jackson), PE Mouse Anti-Human CD19 Clone HIB19 (BD biosciences), FITC Mouse Anti-Human CD20 Clone 2H7 (BD biosciences), BV605 Mouse Anti-Human CD16 Clone 3G8 (BD biosciences), PerCP anti-human CD45 Antibody Clone HI30 (Biolegend), Live/dead aqua dead cell stain (Thermofisher).

### Autonomous growth assessment

To evaluate for autonomous NK cell growth, we maintained control ex vivo-expanded NT NK cells (control or *CISH* KO) and iC9/CAR.19/IL-15-transduced NK cells (control or *CISH* KO) in stem cell growth media without stimulation with feeder cells or addition of exogenous cytokines. Cells were cultured for 45 days and counted using trypan blue exclusion every 3 days.

### Activation of suicide gene *in vitro* and validation *in vivo*

The small-molecule dimerizer AP1903 (10 nM) (generously provided by Bellicum Pharmaceuticals, Inc.; Houston, TX, USA), was added to CB-NK cell cultures for 4 h. The elimination of transduced cells was evaluated by Annexin-V/Live dead staining to detect apoptotic and dead cells. The efficacy of the suicide gene was also tested *in vivo* by treating tumor-bearing mice that had received iC9/CAR.19/IL-15-transduced NK cells (control or *CISH* KO) with two doses of AP1903 (50 μg each) intraperitoneally, 2 days apart on days 7 and 9.^9,43^ Two other groups of mice (control and *CISH* KO) served as control without AP1903 treatment. All mice were sacrificed on day 12, and blood and organs (liver, spleen and bone marrow) were collected and processed for flow cytometry to determine the CAR-NK cell fraction and viability.

### Mitochondria confocal imaging

#### Fluorescence staining of organelles

NK cells were first incubated at 37°C for 40 minutes in 1:1 (v/v) solution of live cell staining buffer (Abcam, Cambridge, UK) and RPMI (Corning, Corning, NY, USA) containing final concentrations of 500 nM MitoTracker™ Deep Red FM (Invitrogen™, Carlsbad, CA, California), 250 nM LysoRed (Abcam, Cambridge, UK) and 1 μM Hoechst 33342 (Sigma, St. Louis, MO, USA) for labeling mitochondria, lysosome and nucleus respectively. Cells then were washed with Hanks’ balanced salt solution (Cellgro, Mediatech Inc., Manassas, VA, USA) + 10% HEPES (Corning, Corning, NY, USA) first and complete culture medium RPMI + 10% FBS (Atlanta Biologicals, Norcross, GA) (R10) for the second wash. The same method was used for each population.

#### Confocal microscopy

∼100,000 cells of each population were loaded in individual wells of 96 glass-bottom plate (MatTek Corporation). A Nikon (Minato, Tokyo, Japan) A1/TiE inverted microscope equipped with a 100x, Nikon, Plan Apo Lambda, oil, 1.45 NA objective was used for imaging. 3D images (z-stacks, 0.3 μm steps, ∼40 slices) were taken from different field of views using DAPI, FITC, TXRed and Cy5 channels.

#### Analyzing confocal images

Z-stacks of 16-bit images were extracted for each channel and processed in ImageJ (National Institutes of Health (NIH), USA) using a series of plugins. First, images were segmented for each channel prior to applying a threshold. Next, the 3D Objects Counter plugin was applied to the image to determine mitochondrial and lysosome regions of interest (ROIs). These ROIs were overlaid onto the original image and measurements were collected afterward. Similarly, the 3D Objects Counter plugin was also used on nucleus but using the original image only. Lastly, tracking of single cell movement was done using the TrackMate plugin^63^ in order to filter out unstable cells upon their movement. All measurements were consolidated in R, where mitochondria, lysosome, and nucleus were matched to their corresponding cell.

### Off-target Identification

The GUIDE-seq method was employed for unbiased discovery of off-target editing events.^28^ In this study, HEK293 cells that constitutively express the *S pyogenes* Cas9 nuclease (“HEK293-Cas9” cells) were used as the source of Cas9. Alt-R® gRNA complexes were formed by combining Alt-R tracrRNA and Alt-R crRNA XT at a 1:1 molar ratio. gRNA complexes were delivered by nucleofection using the Amaxa™ Nucleofector™ 96-well Shuttle™ System (Lonza, Basel, Switzerland). For each nucleofection, 3.5 x 10^5^ HEK293-Cas9 cells were washed with 1X PBS, resuspended in 20 μL solution SF (Lonza) and combined with 10 μM gRNA together with 0.5 μM GUIDE-seq dsDNA donor fragment. This mixture was transferred into one well of a Nucleocuvette™ plate (Lonza) and electroporated using protocol 96-DS-150. DNA was extracted 72 hrs after electroporation using the GeneJET Genomic DNA purification kit (Thermo Fisher Scientific). NGS library preparation, sequencing, and operation of the GUIDE-seq software was performed as previously described,^64^ except that the Needleman-Wunch alignment was incorporated.

### Target enrichment via rhAmpSeq for multiplexed PCR

To better quantify editing at off-target sites found identified with GUIDE-seq, we performed multiplex PCR coupled to amplicon NGS, using rhPCR (PCR executed in the presence of RNaseH2)^65^ with blocked-cleavable primers. Primers were designed by an algorithm (developed by IDT Technologies) for primer cross-comparison and selection based on compatibility with other primers in the multiplex. This amplification technology requires that the primer properly hybridize to a target site before amplification. Mismatches between target and primer prevent unblocking, thereby increasing specificity and eliminating primers dimers. This approach enables efficient production of highly multiplex PCR amplicons in a single tube. For these experiments, gRNA complexes were delivered into HEK293-Cas9 cells as previously described or complexed with Alt-R HiFi Cas9 nuclease v3 to form an active ribonucleoprotein complex (RNP) which was then directly nucleofected into HEK293 cells at 2 μM together with 2 μM Alt-R Cas9 Electroporation Enhancer (IDT). DNA was extracted 48 hrs after electroporation using QuickExtract DNA Extraction Solution (Epicentre). Genomic DNA was extracted in the same way from NK cells for off-target studies comparing WT and HiFi Alt-R Cas9 RNP complexes with editing mediated by optimized delivery conditions, as described previously. Locus-specific amplification with rh-primers was performed for 10 cycles followed by a 1.5x SPRI bead clean-up. An indexing round of PCR was performed for 18 cycles to incorporate sample-unique P5 and P7 indexes followed by a 1x SPRI bead clean-up and library quantification by qPCR (IDT). PCR amplicons were sequenced on an Illumina MiSeq instrument (v2 chemistry, 150 bp paired end reads) (Illumina, San Diego, CA, USA). Data were analyzed using a custom-built pipeline. Data were analyzed using a custom-built pipeline. Data were demultiplexed (Picard tools v2.9; https://github.com/broadinstitute/picard); forward and reverse reads were merged into extended amplicons (flash v1.2.11)^52^; reads were aligned against the GRCh38 genomic reference (minimap2 v2.12), and were assigned to targets in the multiplex primer pool (bedtools tags v2.25)^53^. Reads were re-aligned to the target, favoring alignment choices with INDELs near the Cas9 predicted cut site. At each target, editing was calculated as the percentage of total reads containing an INDEL within a 4bp window of the cut site.

### Statistics

A two way Anova test or student t-test were used as appropriate to compare quantitative differences (mean±s.d.) between groups; *P*-values were two-sided and *P*<0.05 was considered significant. For all bioluminescence experiments, intensity signals were summarized as means±s.d. at baseline and at multiple subsequent time points for each group of mice.^62^ Probabilities of survival were calculated using the Kaplan–Meier method. The indicated statistical tests were performed using Prism software (GraphPad version 7.0c). For the confocal microscopy analysis, data sets were analyzed using unpaired t-tailed test. Data present mean ± 95% confidence interval. Images were assembled using ImageJ.

## Acknowledgments

Supported in part by grants from the M.D. Anderson Cancer Center Lymphoma Moonshot, (CPRIT RP160693) and the National Institutes of Health (1 R01 CA211044-01), PO1 (5P01CA148600-03) and Cancer Center Support (CORE) Grant (CA016672) that support the Flow Cytometry and Cellular Imaging Facility and the RNA sequencing core facility at MD Anderson Cancer Center. We acknowledge the support of the DKMS Mechtild Harf Research Grant and the SITC-Amgen Cancer Immunotherapy in Hematologic Malignancies Fellowship Award awarded to Dr. Daher.

## Author contributions

MD, RB, and PPB performed experiments, interpreted and analyzed data. NB performed western blot and seahorse assay experiments, interpreted and analyzed data. EG, NU, ANC, JL, GO, EE, MeK, VN, MB, LB, and XW assisted with experiments. VM, YX, KC and JW assisted with RNA seq analysis. SK and NP assisted with mass cytometry analysis. MSS, GRR, RT, MSM, GK, HL and MAB performed off target effect experiments and analysis and commented on the manuscript. NWF performed pathologic examination and staining of mice tissues. MaK, ES, RC, KC, EL, SoA, SuA, RJS, SL, and LS commented on the manuscript. DM, MM, LNK, NI, MHS, LL, HS, FL, PPB, and YL provided advice on experiments and commented on the manuscript. KR and MD designed and directed the study. KR, MD and LMF wrote the manuscript.

## Data availability

All requests for raw data and materials will be reviewed by MD Anderson Cancer Center to verify if the request is subject to any intellectual property or confidentiality obligations. Any data and materials that can be shared by the corresponding author will be released freely or via a Material Transfer Agreement if deemed necessary.

## References

1. First-Ever CAR T-cell Therapy Approved in U.S. Cancer Discov 2017; 7(10): OF1.

2. FDA Approves Second CAR T-cell Therapy. Cancer Discov 2017.

3. Hartmann J, Schussler-Lenz M, Bondanza A, Buchholz CJ. Clinical development of CAR T cells-challenges and opportunities in translating innovative treatment concepts. EMBO Mol Med 2017; 9(9): 1183–97.

4. Morvan MG, Lanier LL. NK cells and cancer: you can teach innate cells new tricks. Nat Rev Cancer 2016; 16(1): 7–19.

5. Simonetta F, Alvarez M, Negrin RS. Natural Killer Cells in Graft-versus-Host-Disease after Allogeneic Hematopoietic Cell Transplantation. Front Immunol 2017; 8: 465.

6. Mehta RS, Shpall EJ, Rezvani K. Cord Blood as a Source of Natural Killer Cells. Front Med (Lausanne) 2015; 2: 93.

7. Sarvaria A, Jawdat D, Madrigal JA, Saudemont A. Umbilical Cord Blood Natural Killer Cells, Their Characteristics, and Potential Clinical Applications. Front Immunol 2017; 8: 329.

8. Daher M, Rezvani K. Next generation natural killer cells for cancer immunotherapy: the promise of genetic engineering. Curr Opin Immunol 2018; 51: 146–53.

9. Liu E, Tong Y, Dotti G, et al. Cord blood NK cells engineered to express IL-15 and a CD19-targeted CAR show long-term persistence and potent antitumor activity. Leukemia 2017.

10. Sharma P, Allison JP. Immune checkpoint targeting in cancer therapy: toward combination strategies with curative potential. Cell 2015; 161(2): 205–14.

11. Sharma P, Allison JP. The future of immune checkpoint therapy. Science 2015; 348(6230): 56–61.

12. Davis ZB, Vallera DA, Miller JS, Felices M. Natural killer cells unleashed: Checkpoint receptor blockade and BiKE/TriKE utilization in NK-mediated anti-tumor immunotherapy. Semin Immunol 2017; 31: 64–75.

13. Krebs DL, Hilton DJ. SOCS proteins: negative regulators of cytokine signaling. Stem Cells 2001; 19(5): 378–87.

14. Linossi EM, Babon JJ, Hilton DJ, Nicholson SE. Suppression of cytokine signaling: the SOCS perspective. Cytokine Growth Factor Rev 2013; 24(3): 241–8.

15. Yoshimura A, Nishinakamura H, Matsumura Y, Hanada T. Negative regulation of cytokine signaling and immune responses by SOCS proteins. Arthritis Res Ther 2005; 7(3): 100–10.

16. Inagaki-Ohara K, Hanada T, Yoshimura A. Negative regulation of cytokine signaling and inflammatory diseases. Curr Opin Pharmacol 2003; 3(4): 435–42.

17. Delconte RB, Kolesnik TB, Dagley LF, et al. CIS is a potent checkpoint in NK cell-mediated tumor immunity. Nat Immunol 2016; 17(7): 816–24.

18. Cherkassky L, Morello A, Villena-Vargas J, et al. Human CAR T cells with cell-intrinsic PD-1 checkpoint blockade resist tumor-mediated inhibition. J Clin Invest 2016; 126(8): 3130–44.

19. Chong EA, Melenhorst JJ, Lacey SF, et al. PD-1 blockade modulates chimeric antigen receptor (CAR)-modified T cells: refueling the CAR. Blood 2017; 129(8): 1039–41.

20. Mukherjee M, Mace EM, Carisey AF, Ahmed N, Orange JS. Quantitative Imaging Approaches to Study the CAR Immunological Synapse. Mol Ther 2017; 25(8): 1757–68.

21. Putz EM, Guillerey C, Kos K, et al. Targeting cytokine signaling checkpoint CIS activates NK cells to protect from tumor initiation and metastasis. Oncoimmunology 2017; 6(2): e1267892.

22. Felices M, Lenvik AJ, McElmurry R, et al. Continuous treatment with IL-15 exhausts human NK cells via a metabolic defect. JCI Insight 2018; 3(3).

23. Conlon KC, Lugli E, Welles HC, et al. Redistribution, hyperproliferation, activation of natural killer cells and CD8 T cells, and cytokine production during first-in-human clinical trial of recombinant human interleukin-15 in patients with cancer. J Clin Oncol 2015; 33(1): 74–82.

24. Waldmann TA, Lugli E, Roederer M, et al. Safety (toxicity), pharmacokinetics, immunogenicity, and impact on elements of the normal immune system of recombinant human IL-15 in rhesus macaques. Blood 2011; 117(18): 4787–95.

25. Donnelly RP, Loftus RM, Keating SE, et al. mTORC1-dependent metabolic reprogramming is a prerequisite for NK cell effector function. J Immunol 2014; 193(9): 4477–84.

26. Marcais A, Cherfils-Vicini J, Viant C, et al. The metabolic checkpoint kinase mTOR is essential for IL-15 signaling during the development and activation of NK cells. Nat Immunol 2014; 15(8): 749–57.

27. Donnelly RP, Finlay DK. Glucose, glycolysis and lymphocyte responses. Mol Immunol 2015; 68(2 Pt C): 513–9.

28. Tsai SQ, Zheng Z, Nguyen NT, et al. GUIDE-seq enables genome-wide profiling of off-target cleavage by CRISPR-Cas nucleases. Nat Biotechnol 2015; 33(2): 187–97.

29. Vakulskas CA, Dever DP, Rettig GR, et al. A high-fidelity Cas9 mutant delivered as a ribonucleoprotein complex enables efficient gene editing in human hematopoietic stem and progenitor cells. Nat Med 2018; 24(8): 1216–24.

30. Lanuza PM, Pesini C, Arias MA, Calvo C, Ramirez-Labrada A, Pardo J. Recalling the Biological Significance of Immune Checkpoints on NK Cells: A Chance to Overcome LAG3, PD1, and CTLA4 Inhibitory Pathways by Adoptive NK Cell Transfer? Front Immunol 2019; 10: 3010.

31. Pesce S, Greppi M, Tabellini G, et al. Identification of a subset of human natural killer cells expressing high levels of programmed death 1: A phenotypic and functional characterization. J Allergy Clin Immunol 2017; 139(1): 335–46 e3.

32. Concha-Benavente F, Kansy B, Moskovitz J, Moy J, Chandran U, Ferris RL. PD-L1 Mediates Dysfunction in Activated PD-1(+) NK Cells in Head and Neck Cancer Patients. Cancer Immunol Res 2018; 6(12): 1548–60.

33. Benson DM, Jr., Bakan CE, Mishra A, et al. The PD-1/PD-L1 axis modulates the natural killer cell versus multiple myeloma effect: a therapeutic target for CT-011, a novel monoclonal anti-PD-1 antibody. Blood 2010; 116(13): 2286–94.

34. Liu Y, Cheng Y, Xu Y, et al. Increased expression of programmed cell death protein 1 on NK cells inhibits NK-cell-mediated anti-tumor function and indicates poor prognosis in digestive cancers. Oncogene 2017; 36(44): 6143–53.

35. Jones RG, Pearce EJ. MenTORing Immunity: mTOR Signaling in the Development and Function of Tissue-Resident Immune Cells. Immunity 2017; 46(5): 730–42.

36. Weichhart T, Hengstschlager M, Linke M. Regulation of innate immune cell function by mTOR. Nat Rev Immunol 2015; 15(10): 599–614.

37. Preston GC, Sinclair LV, Kaskar A, et al. Single cell tuning of Myc expression by antigen receptor signal strength and interleukin-2 in T lymphocytes. EMBO J 2015; 34(15): 2008–24.

38. Cichocki F, Hanson RJ, Lenvik T, et al. The transcription factor c-Myc enhances KIR gene transcription through direct binding to an upstream distal promoter element. Blood 2009; 113(14): 3245–53.

39. O’Brien KL, Finlay DK. Immunometabolism and natural killer cell responses. Nat Rev Immunol 2019; 19(5): 282–90.

40. Gardiner CM, Finlay DK. What Fuels Natural Killers? Metabolism and NK Cell Responses. Front Immunol 2017; 8: 367.

41. Papa S, Choy PM, Bubici C. The ERK and JNK pathways in the regulation of metabolic reprogramming. Oncogene 2019; 38(13): 2223–40.

42. Sukumar M, Liu J, Ji Y, et al. Inhibiting glycolytic metabolism enhances CD8+ T cell memory and antitumor function. J Clin Invest 2013; 123(10): 4479–88.

43. Hoyos V, Savoldo B, Quintarelli C, et al. Engineering CD19-specific T lymphocytes with interleukin-15 and a suicide gene to enhance their anti-lymphoma/leukemia effects and safety. Leukemia 2010; 24(6): 1160–70.

44. Vera J, Savoldo B, Vigouroux S, et al. T lymphocytes redirected against the kappa light chain of human immunoglobulin efficiently kill mature B lymphocyte-derived malignant cells. Blood 2006; 108(12): 3890–7.

45. Moreno-Mateos MA, Vejnar CE, Beaudoin JD, et al. CRISPRscan: designing highly efficient sgRNAs for CRISPR-Cas9 targeting in vivo. Nat Methods 2015; 12(10): 982–8.

46. Li D, Qiu Z, Shao Y, et al. Heritable gene targeting in the mouse and rat using a CRISPR-Cas system. Nat Biotechnol 2013; 31(8): 681–3.

47. Gundry MC, Brunetti L, Lin A, et al. Highly Efficient Genome Editing of Murine and Human Hematopoietic Progenitor Cells by CRISPR/Cas9. Cell Rep 2016; 17(5): 1453–61.

48. Kolesnik TB, Nicholson SE. Analysis of Suppressor of Cytokine Signalling (SOCS) gene expression by real-time quantitative PCR. Methods Mol Biol 2013; 967: 235–48.

49. Rouce RH, Shaim H, Sekine T, et al. The TGF-beta/SMAD pathway is an important mechanism for NK cell immune evasion in childhood B-acute lymphoblastic leukemia. Leukemia 2016; 30(4): 800–11.

50. Banerjee PP, Orange JS. Quantitative measurement of F-actin accumulation at the NK cell immunological synapse. J Immunol Methods 2010; 355(1-2): 1–13.

51. Sanborn KB, Rak GD, Mentlik AN, Banerjee PP, Orange JS. Analysis of the NK cell immunological synapse. Methods Mol Biol 2010; 612: 127–48.

52. Muftuoglu M, Olson A, Marin D, et al. Allogeneic BK Virus-Specific T Cells for Progressive Multifocal Leukoencephalopathy. N Engl J Med 2018; 379(15): 1443–51.

53. Van Gassen S, Callebaut B, Van Helden MJ, et al. FlowSOM: Using self-organizing maps for visualization and interpretation of cytometry data. Cytometry A 2015; 87(7): 636–45.

54. Kordasti S, Costantini B, Seidl T, et al. Deep-phenotyping of Tregs identifies an immune signature for idiopathic aplastic anemia and predicts response to treatment. Blood 2016.

55. Povoleri GAM, Nova-Lamperti E, Scotta C, et al. Human retinoic acid-regulated CD161(+) regulatory T cells support wound repair in intestinal mucosa. Nat Immunol 2018; 19(12): 1403–14.

56. Vivian J, Rao AA, Nothaft FA, et al. Toil enables reproducible, open source, big biomedical data analyses. Nat Biotechnol 2017; 35(4): 314–6.

57. Love MI, Huber W, Anders S. Moderated estimation of fold change and dispersion for RNA-seq data with DESeq2. Genome Biol 2014; 15(12): 550.

58. Luo W, Friedman MS, Shedden K, Hankenson KD, Woolf PJ. GAGE: generally applicable gene set enrichment for pathway analysis. BMC Bioinformatics 2009; 10: 161.

59. Barbie DA, Tamayo P, Boehm JS, et al. Systematic RNA interference reveals that oncogenic KRAS-driven cancers require TBK1. Nature 2009; 462(7269): 108–12.

60. Hanzelmann S, Castelo R, Guinney J. GSVA: gene set variation analysis for microarray and RNA-seq data. BMC Bioinformatics 2013; 14: 7.

61. Liberzon A, Birger C, Thorvaldsdottir H, Ghandi M, Mesirov JP, Tamayo P. The Molecular Signatures Database (MSigDB) hallmark gene set collection. Cell Syst 2015; 1(6): 417–25.

62. Shah N, Martin-Antonio B, Yang H, et al. Antigen presenting cell-mediated expansion of human umbilical cord blood yields log-scale expansion of natural killer cells with anti-myeloma activity. PLoS One 2013; 8(10): e76781.

63. Tinevez JY, Perry N, Schindelin J, et al. TrackMate: An open and extensible platform for single-particle tracking. Methods 2017; 115: 80–90.

64. Tsai SQ, Topkar VV, Joung JK, Aryee MJ. Open-source guideseq software for analysis of GUIDE-seq data. Nat Biotechnol 2016; 34(5): 483.

65. Dobosy JR, Rose SD, Beltz KR, et al. RNase H-dependent PCR (rhPCR): improved specificity and single nucleotide polymorphism detection using blocked cleavable primers. BMC Biotechnol 2011; 11: 80.

