## Supplementary Materials for "CIS checkpoint deletion enhances the fitness of cord blood derived natural killer cells transduced with a chimeric antigen receptor"

**Supplementary Figures**

Supplementary Figure 1

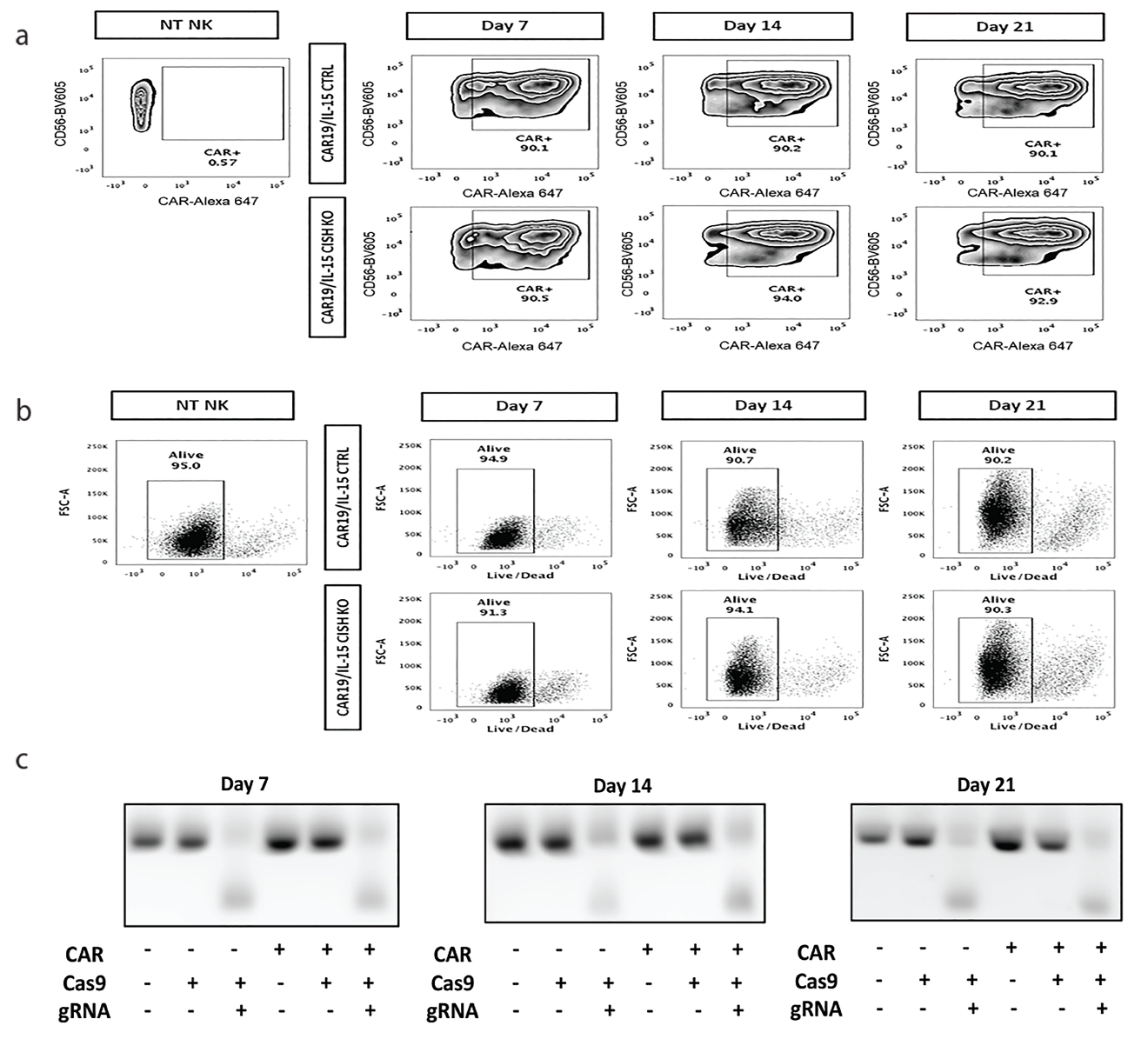

Supplementary Figure 2

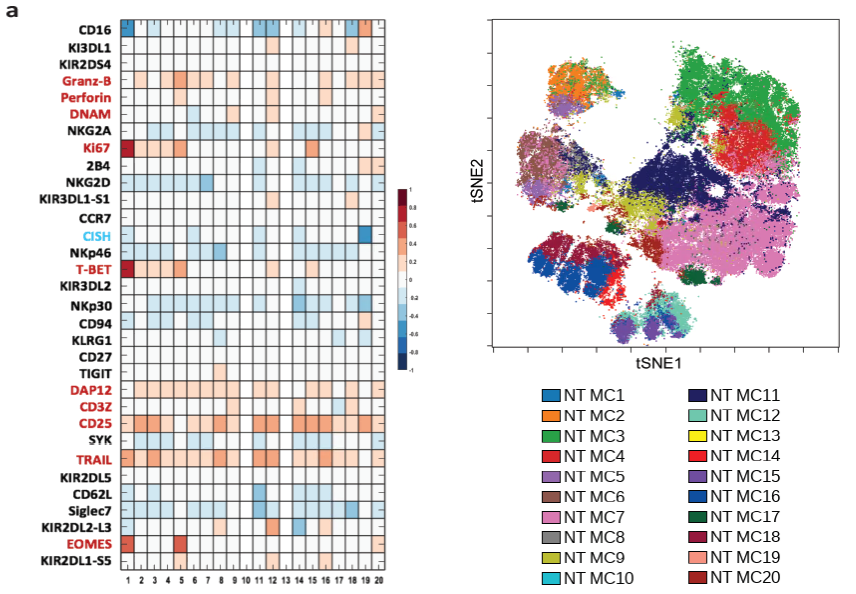

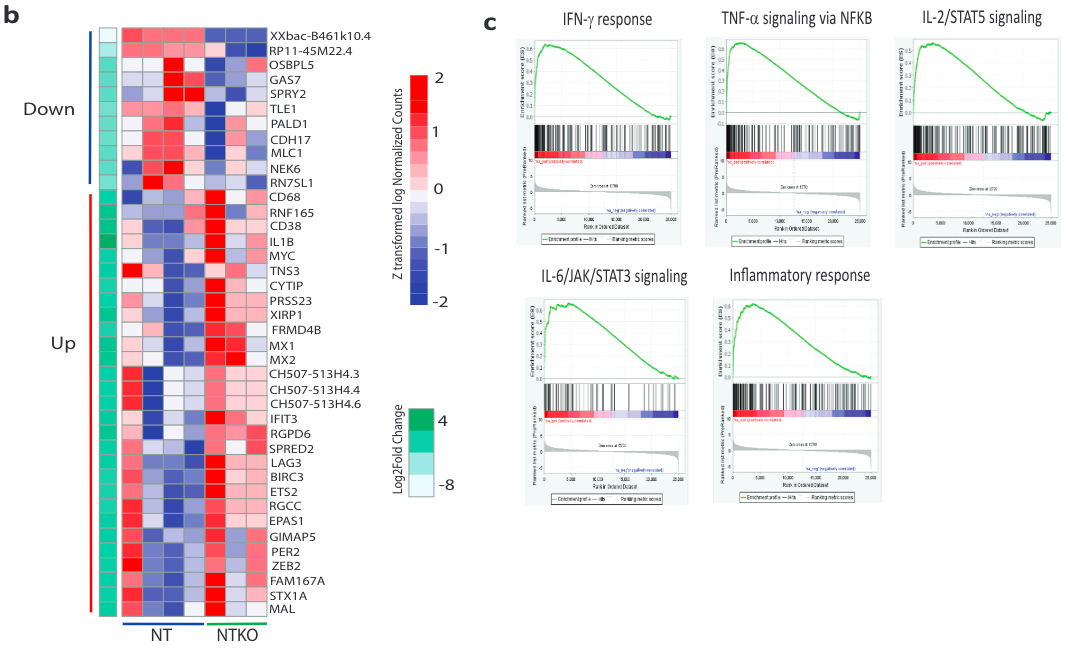

Supplementary Figure 3

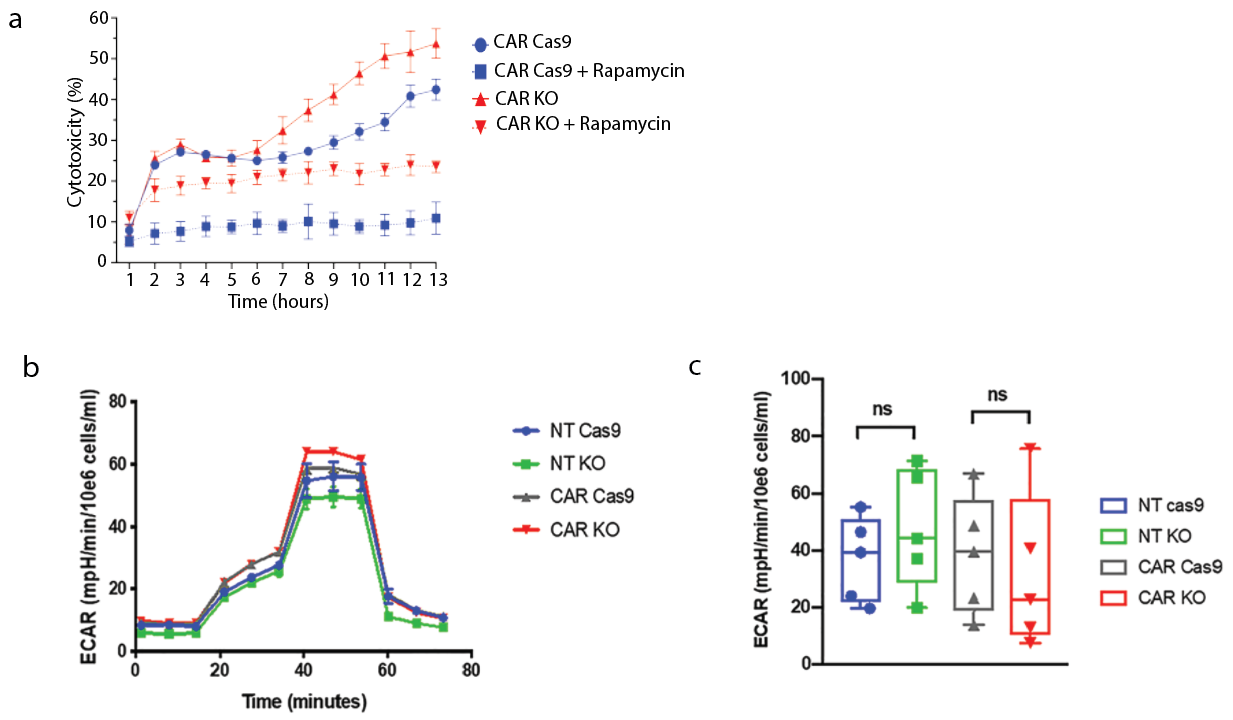

Supplementary Figure 4

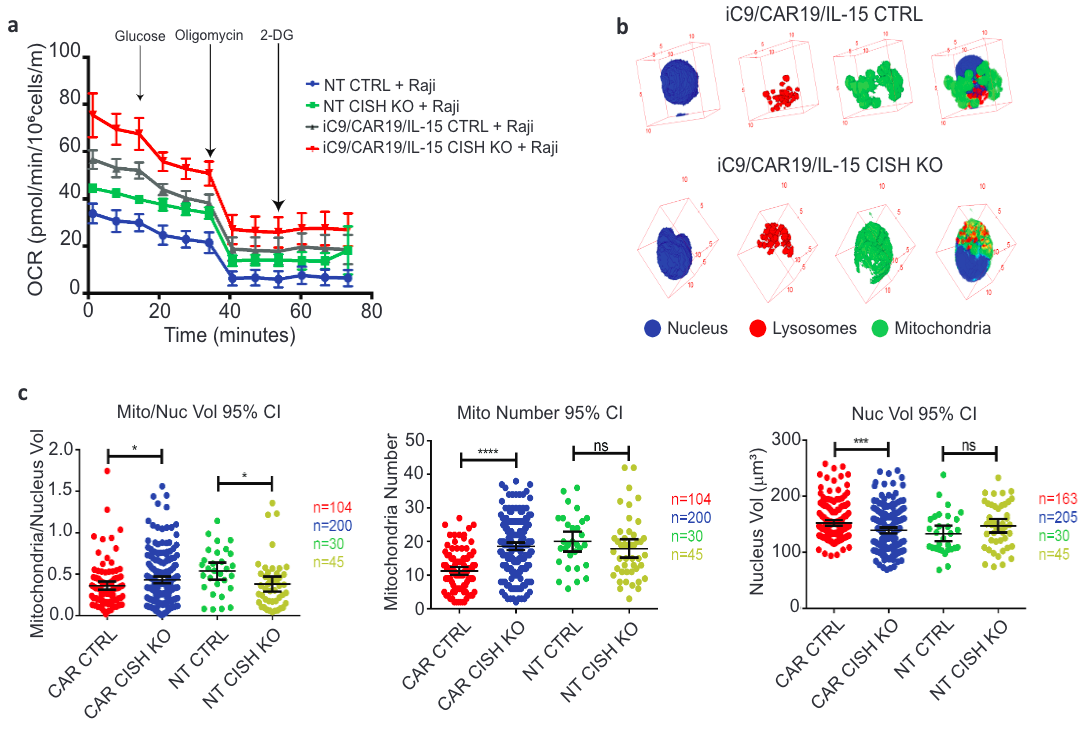

Supplementary Figure 5

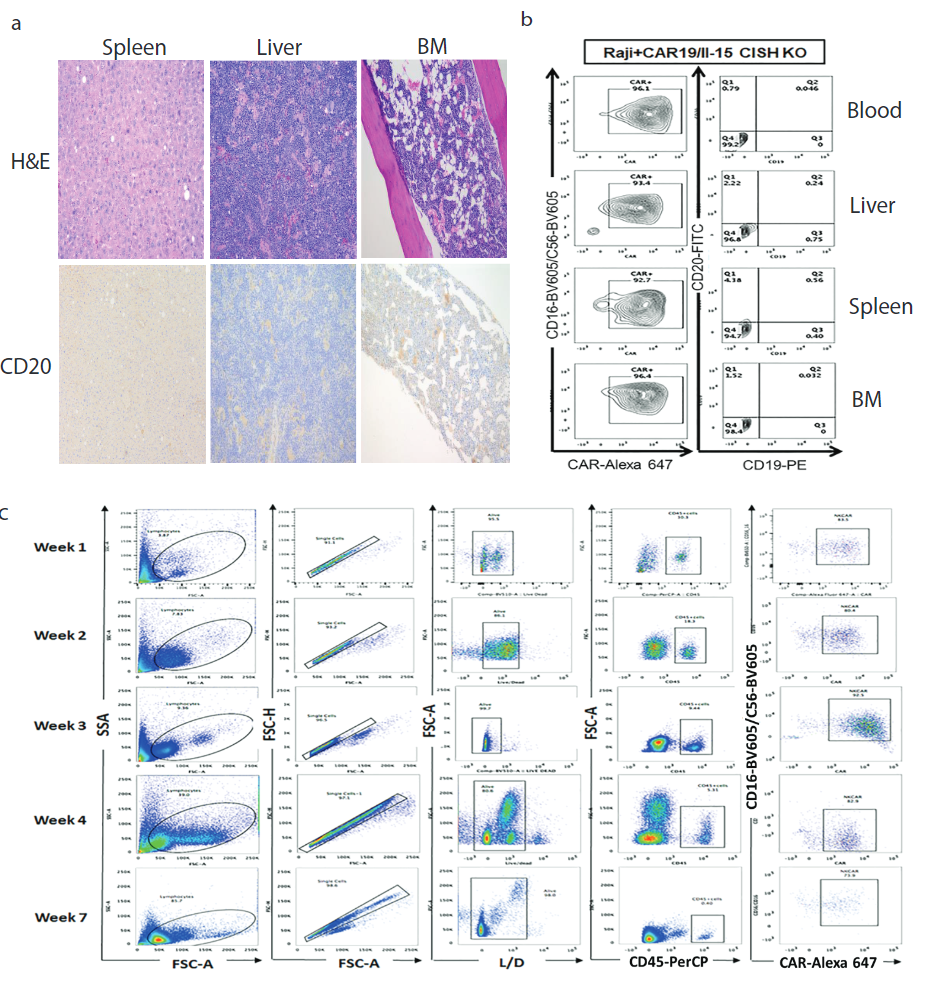

Supplementary Figure 6

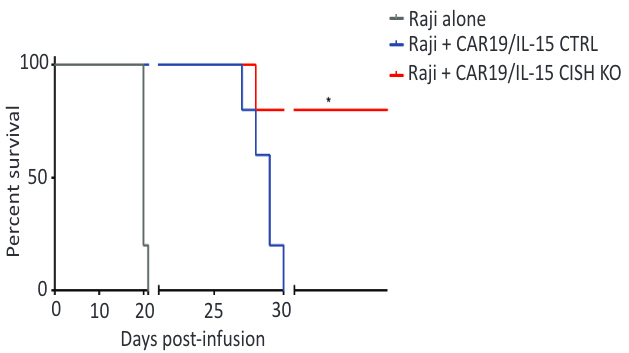

Supplemetary Figure 7

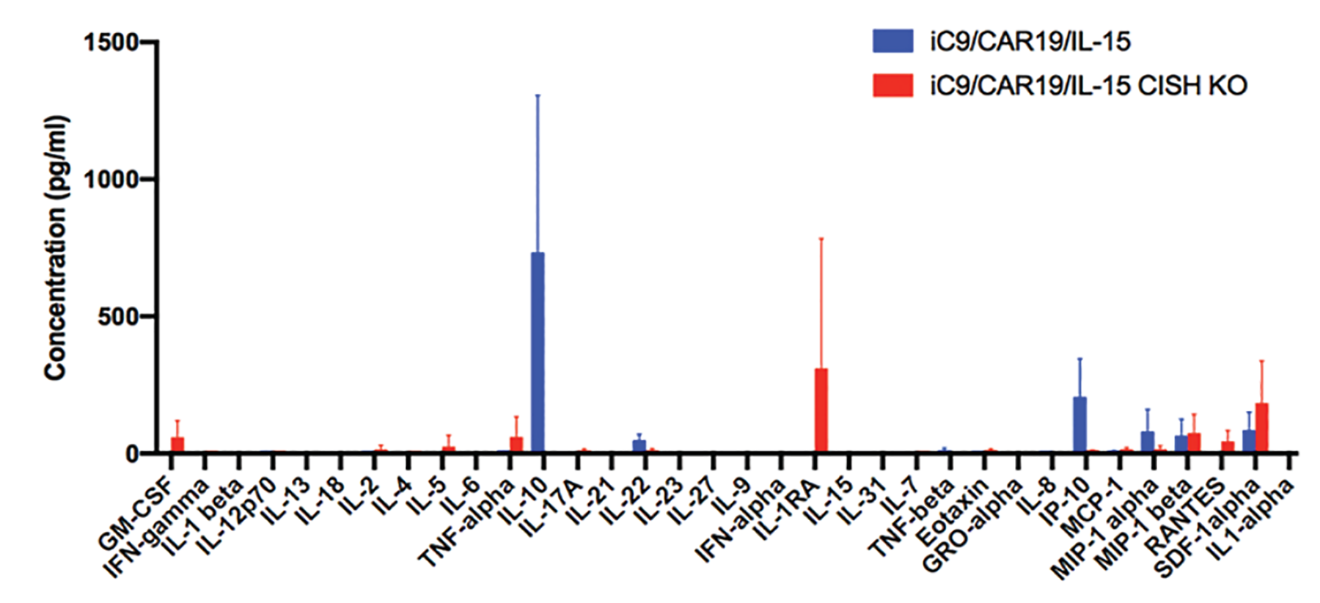

Supplementary Figure 8

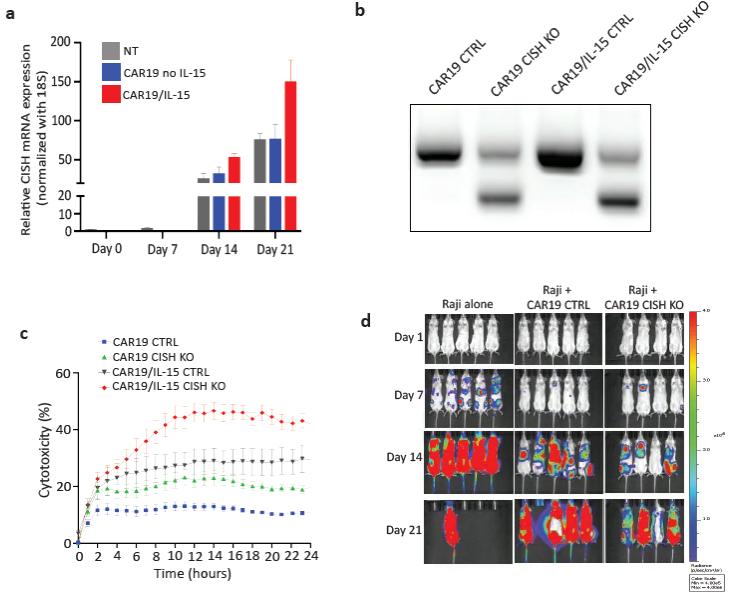

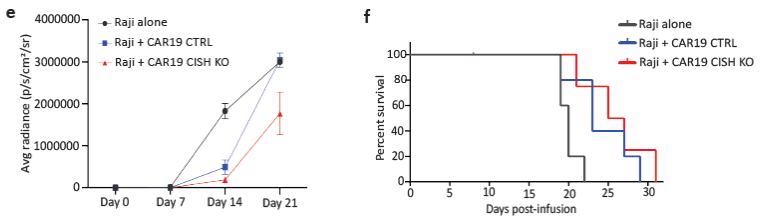

Supplementary Figure 9

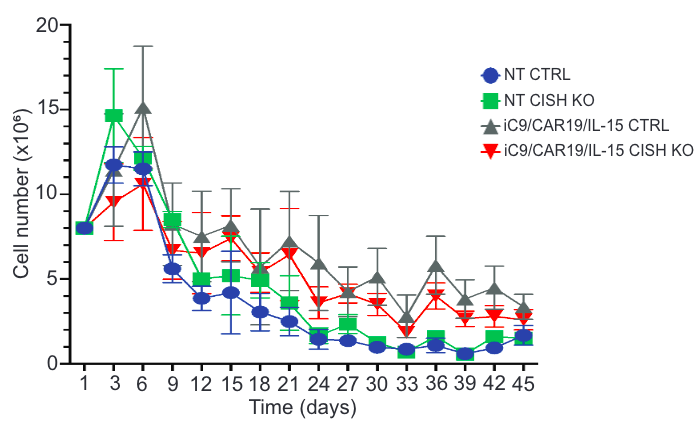

Supplementary Figure 10

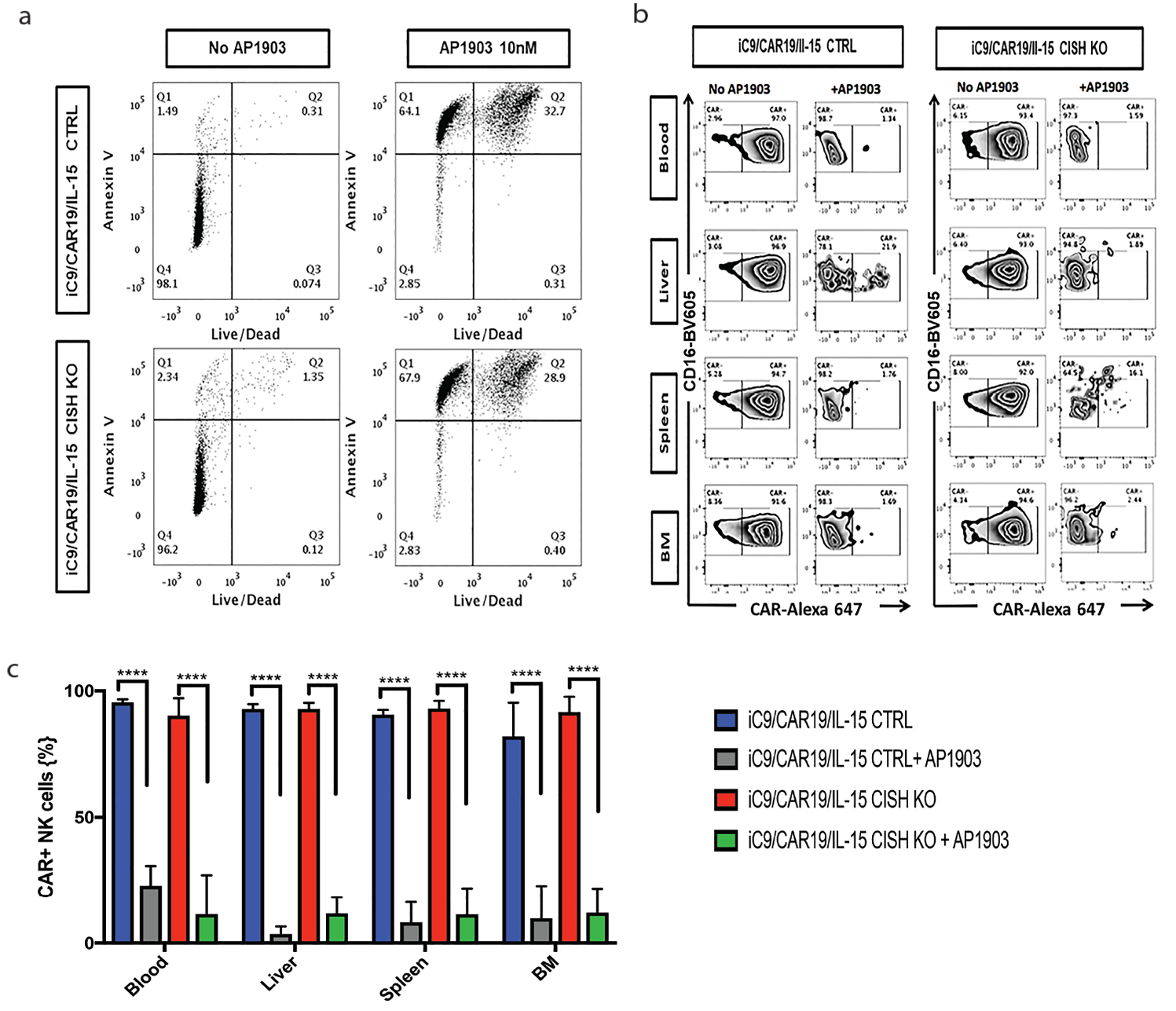

**Supplementary Figure Legends**

**Supplementary Figure 1. iC9/CAR19/IL-15 transduction, viability and *CISH* KO efficiency are stable over time.** CAR expression (**a**) and viability of (**b**) of iC9/CAR19/IL-15 transduced NK cells vs. *CISH* KO iC9/CAR19/IL-15 transduced NK cells over time. Inset values indicate the percentage of CAR+ NK cells (**a**) and alive cells (**b**) from each group. **c**, The *CISH* KO efficiency in NT and iC9/CAR19/IL-15-expressing NK cells at days 7, 14 and 21 following CRISPR/Cas9-mediated knockout was determined by PCR analysis.

**Supplementary Figure 2. Molecular signature of *CISH* KO NT NK cells. a,** Comparative heatmap of mass cytometry data shows expression of transcription factors and cytotoxicity markers in NT *CISH* KO compared to NT CTRL NK cells. Each column represents a separate cluster identified by FlowSOM analysis and each row reflects the expression of a certain marker for each annotation. Color scale shows the expression level of each marker, with red representing higher expression and blue lower expression of NT *CISH* KO NK cells. The t-SNE map generated from FlowSOM analysis in the right panel shows the 20 NK cells metaclusters (MC) represented in the mass cytometry heat map in the left panel. **b**, Heat map displays RNA sequencing data of genes that were differentially expressed (q < 0.1 and absolute log2foldchange > 0.8) in NT *CISH* KO compared to NT CTRL NK cells (n=2). **c**, Gene set enrichment analysis (GSEA) plots showing enrichment in IFN-γ response, TNF-α signaling via NF-kB, IL-16/JAK/STAT3, IL-2/STAT5 signaling and inflammatory response in NT *CISH* KO compared to NT CTRL NK cells.

**Supplementary Figure 3. The mTORC1 pathway and the presence of tumor are important for the optimal effect of *CISH* KO on the activiy of iC9/CAR19/IL-15 NK cells. a,** Cytotoxicity of iC9/CAR19/IL-15 *CISH* KO NK cells and iC9/CAR19/IL-15 control (Cas9) NK cells treated with or without rapamycin (100ng/ml) over 24 hours at 1:1 E:T ratio was measured by Incucyte live imaging cell killing assay (n=2). Bars represent mean values with standard deviation. Solid lines represent untreated cells and dotted lines represent cells treated with rapamycin. Red lines represent iC9/CAR19/IL-15 *CISH* KO NK cells and blue lines represent iC9/CAR19/IL-15 control (Cas9) NK cell. **b**, A series of extracellular acidification rate (ECAR) were calculated for NT CTRL (blue lines), NT *CISH* KO (green lines), iC9/CAR19/IL-15 CTRL (grey lines) or iC9/CAR19/IL-15 *CISH* KO (red lines) NK cells cultured alone and subsequently treated with 2 g/L D-glucose, 1 μM oligomycin and 100 mM 2-Deoxyglucose (2-DG). A representative graph is shown from five independent experiments. **c**, Box plots summarize the ECAR data by NT CTRL (blue box), NT *CISH* KO (green box), iC9/CAR19/IL-15 CTRL (grey box) or iC9/CAR19/IL-15 *CISH* KO (red box) NK cells cultured alone (n=5). NS, not statistical significance.

**Supplementary Fig. 4. *CISH* KO CAR19/IL-15 NK cells have increased mitochondrial number, volume and activity.** **a**, Oxygen consumption rates (OCR) were calculated for NT CTRL (blue lines), *CISH* KO NT (green lines), iC9/CAR19/IL-15 CTRL (grey lines) or *CISH* KO iC9/CAR19/IL-15 (red lines) NK cells co-cultured with Raji targets for 2 hrs and subsequently purified and treated with 2 g/L D-glucose, 1 μM oligomycin and 100 mM 2-Deoxyglucose (2-DG). A representative graph is shown from 2 independent experiments. Bars represent mean values with standard deviation. **b**, Spatial representation of confocal images of nucleus (blue), lysosomes (red) and mitochondria (green) in both iC9/CAR19/IL-15 CTRL and *CISH* KO iC9/CAR19/IL-15. **c**, Dot plot graphs representing mitochondrial/nuclear volume (Mito/Nuc Vol) ratio, mitochondrial number and nuclear volume (Nuc Vol) from iC9/CAR19/IL-15 CTRL (red dots), iC9/CAR19/IL-15 *CISH* KO (blue dots), NT CTRL (green dots) or NT *CISH* KO (yellow dots) NK cells.

**Supplementary Figure 5. No evidence of Raji lymphoma in NSG mice receiving 10 x 10^6^ iC9/CAR19/IL-15 *CISH* KO NK cells. a,** NSG mice >1 year after treatment with Raji plus one dose of 10 x 10^6^ iC9/CAR19/IL-15 *CISH* KO NK cells were sacrificed and examined for dysregulated NK cell growth or leukemia/lymphoma. Photomicrographs of spleen (left), liver (center) or bone marrow (BM; right) show no evidence of pathology. H&E staining (top panels; magnification 100x) and CD20 immunohistochemical staining (lower panels; magnification 100x). The micrographs are from a representative NSG mouse treated with iC9/CAR19/IL-15 *CISH* KO. **b**, Flow cytometry data showing the presence of CAR+ NK cells and the absence of Raji lymphoma (CD19+CD20+) in blood, spleen, liver and bone marrow (BM) of an NSG mouse treated with Raji plus one dose of 10 x 10^6^ iC9/CAR19/IL-15 *CISH* KO NK cells and sacrificed on day 54. **c**, Flow plots showing the expression levels of NK cells in mice receiving *CISH* KO iC9/CAR19/IL-15 cells vs. iC9/CAR19/IL-15 NK cell controls up to 7 weeks after initial infusion. Inset values indicate the frequency of positive cells from each group.

**Supplementary Figure 6. *CISH* KO iC9/CAR19/IL-15 NK cells improve survival in a Raji lymphoma mouse model.** Survival curve of an independent mouse experiment where mice received either Raji alone (n=5, grey) or Raji plus 1 x 10^7^ CAR19/IL-15 (n=5, blue) or Raji plus 1 x 10^7^ CAR19/IL-15 *CISH* KO (n=5, red). Statistical significance is represented by *p≤ 0.05 for the comparison of the red and blue curves.

**Supplementary Figure 7. Expression levels of inflammatory cytokines following infusion of *CISH* KO or control iC9/CAR19/IL-15 NK cells in mice engrafted with Raji lymphoma.** Bar graphs showing the production of 34 inflammatory cytokines in the plasma of mice 14 days following iC9/CAR19/IL-15 CTLT (blue) vs. iC9/CAR19/IL-15 *CISH* KO (red) NK cell infusion as measured by ProcartaPlex Immunoassay for human cytokines (n=5 in each group). Bars represent mean values with standard deviation.

**Supplementary Figure 8. *CISH* KO in CAR19 NK cells (without IL-15) does not lead to improvement in the anti-tumor activity of CAR19 NK cells.** **a**, Bar graphs showing the relative mRNA expression levels of *CISH* on days 0, 7, 14 and 21 of expansion from NT (grey) CAR19-transduced (without IL-15; blue) and iC9/CAR19/IL-15-transduced (red) NK cells by reverse transcription polymerase chain reaction (RT-PCR) (n=2). Note that on days 0 and 7 only data for NT NK cells are included since the CAR transduction step was performed on day 4 of expansion. 18 S ribosomal RNA (18S) was used as the internal reference gene. Bars represent mean values with standard deviation. **b**, The *CISH* KO efficiency was determined by PCR for CAR19 CTRL, CAR19 *CISH* KO, CAR19/IL-15 CTRL and CAR19/IL-15 *CISH* KO NK cells. **c**, Cytotoxicity of CAR19 CTRL (blue lines), CAR19 *CISH* KO (green lines), iC9/CAR19/IL-15 CTRL (grey lines) or iC9/CAR19/IL-15 *CISH* KO (red lines) NK cells against Raji targets over 24 hours at 1:1 E:T ratio as measured by Incucyte live imaging cell killing assay (n=3). Bars represent mean values with standard deviation. **d,** BLI data from three groups of NSG mice treated with Raji alone (n=5), Raji plus one dose of 10 x 10^6^ CAR19 CTRL NK cells (n=5) or CAR19 *CISH* KO NK cells (n=4). **e,** The average radiance and **f,** survival curves are shown for the three groups of mice described in panel d.

**Supplementary Figure 9. Autonomous growth study for engineered NK cells.** Curve graphs representing the autonomous growth of NT CTRL (blue), NT *CISH* KO (green), CAR19/IL-15 CTRL (grey) and CAR19/Il-15 (red) *CISH* KO NK cells cultured in media without the addition of exogenous cytokines or feeder cells for 45 days from 3 independent experiments. Bars represent mean values with standard deviation.

**Supplementary Figure 10. Activation of inducible caspase-9 gene eliminates iC9/CAR19/IL-15 CTRL and iC9/CAR19/IL-15 *CISH* KO NK cells. a,** Addition of AP1903 (10nM) to cultures of iC9/CAR19/IL-15 CTRL or iC9/CAR19/IL-15 *CISH* KO NK cells induces cell apoptosis within 4 hours as assessed by annexin-V-7AAD staining. Data are representative of 3 independent experiments. **b**, NSG mice engrafted with Raji cells and infused with 10 x 10^6^ iC9/CAR19/IL-15 CTRL or iC9/CAR19/IL-15 *CISH* KO NK cells were either followed without intervention or treated 7 days later with 2 doses of the AP1903 dimerizer (50μg) i.p two days apart (n=5 mice per group). FACS plots from the blood and tissue of one representative mouse from each group are shown. In the mice that received AP1903 injections, iC9/CAR19/IL-15 NK cells (CTRL or *CISH* KO) were significantly reduced in all organs harvested 3 days after the last dose of AP1903 as measured by the frequencies of CAR-positive NK cells in blood, liver, spleen and bone marrow (BM) by flow cytometry. **c**, Bar graphs showing the percentage of CAR+ NK cells in blood, liver, spleen and bone marrow (BM) specimen collected from NSG mice engrafted with Raji cells and infused with 10 x 10^6^ iC9/CAR19/IL-15 CTRL or iC9/CAR19/IL-15 *CISH* KO NK cells and treated with or without AP1903 dimerizer (50μg i.p x2 doses). Bars represent mean values with standard deviation, ****p ≤ 0.0001.

**Supplementary Table 1. List of antibodies used for mass cytometry panel**

|  | TARGET | Clone | ISOTOPE | Source |
| --- | --- | --- | --- | --- |
| 1 | CD45 | HI30 | 89Y | Fluidigm |
| 2 | CD57 | HCD57 | 115In | Biolegend |
| 3 | KIR2DL1/S5 | HP-MA4 | 141Pr | Biolegend |
| 4 | EOMES | WD1928 | 142Nd | Thermo Fisher |
| 5 | KIR2DL2/L3 | DX27 | 143Nd | Biolegend |
| 6 | Siglec 7 | EMR8-5 | 144Nd | BD Biosciences |
| 7 | CD62L | DREG-56 | 145Nd | Biolegend |
| 8 | KIR2DL5 | UP-R1 | 146Nd | Miltenyi |
| 9 | CD20 | 2H7 | 147Sm | Fluidigm |
| 10 | TRAIL | RIK-2 | 148Nd | BD Bioscience |
| 11 | SYK | 4D10.2 | 149Sm | Biolegend |
| 12 | KIR2DL4 | 181703 | 150Nd | R&D |
| 13 | CD25 | 2A3 | 151Eu | Miltenyi |
| 14 | CD3Z | 6B10.2 | 152Sm | Biolegend |
| 15 | DAP12 | 406288 | 153Eu | R&D |
| 16 | TIGIT | MBSA43 | 154Sm | Thermo Fisher |
| 17 | CD27 | L128 | 155Gd | Bd Bioscience |
| 18 | KLRG1 | 13F12F2 | 156Gd | Thermo Fisher |
| 19 | CD94 | DX22 | 158Gd | Biolegend |
| 20 | NKP30 | Z5 | 159Tb | Fluidigm |
| 21 | KIR3DL2 | 539304 | 160Gd | R&D |
| 22 | T-BET | 4B10 | 161Dy | Biolegend |
| 23 | NKP46 | BAB281 | 162Dy | Fluidigm |
| 24 | CISH | Polyclonal | 163Dy | R&D |
| 25 | CCR7 | G043H7 | 164Dy | Biolegend |
| 26 | NKG2D | ON72 | 166Er | Beckman Coulter |
| 27 | 2B4 | C1.7 | 167Er | Thermo Fisher |
| 28 | KI67 | Ki67 | 168Er | Biolegend |
| 29 | NKG2A | Z199 | 169Tb | Fluidigm |
| 30 | CD3 | UCHT-1 | 170Er | Biolegend |
| 31 | DNAM | DX11 | 171Yb | BD Bioscience |
| 32 | Perforin | dG9 | 172Yb | Biolegend |
| 33 | Granzyme B | GB11 | 173Yb | BD Bioscience |
| 34 | KIR2DS4 | JJC11.6 | 174Yb | Miltenyi |
| 35 | KIR3DL1 | DX9 | 175Lu | BD Bioscience |
| 36 | CD56 | NCAM16.2 | 176Yb | BD Bioscience |
| 37 | CD16 | 3G8 | 209Bi | Fluidigm |
|  | Cisplatin |  | 198Pt | Fluidigm |
